# Wheat Enhanced Disease Resistance EMS-Mutants Include Lesion-mimics With Adult Plant Resistance to Stripe Rust

**DOI:** 10.1101/2024.05.10.593581

**Authors:** China Lunde, Kyungyong Seong, Rakesh Kumar, Andrew Deatker, Bhavit Chhabra, Meinan Wang, Shivreet Kaur, Sarah Song, Ann Palayur, Cole Davies, William Cumberlich, Upinder Gill, Nidhi Rawat, Xianming Chen, Meriem Aoun, Christopher Mundt, Ksenia V Krasileva

## Abstract

Tetraploid durum wheat *Triticum turgidum* subsp. *durum* cv ‘Kronos’ has extensive genetic variation resources, including a sequenced and cataloged ethyl methanesulfonate (EMS) mutagenized population. To utilize this allelic diversity, we screened over 2,000 mutant lines and identified over 30 enhanced disease resistance (EDR) mutants in a forward genetic field screen against stripe rust. Sixteen of the EDR lines have persistent resistance to stripe rust after four years, and several mutants showed differential disease responses against other fungal pathogens, indicating that the lines possess diverse alleles that affect multiple routes of pathogen suppression. Five of these 16 lines showed spontaneous lesion formation in the absence of pathogens. Only one showed a reduction in kernel weight under pathogen pressure, a testimony to the high mutational density that wheat can tolerate. Phenotypic selection for resistance at the adult stage identified useful EMS alleles for stripe rust resistance. The mutations in the 16 EDR lines were newly mapped to a recently released long-read Kronos genome to enhance their utility in molecular breeding for fungal resistance and for fundamental studies of plant-pathogen interactions.

## Introduction

Stripe (yellow) rust of wheat, caused by the obligate biotrophic fungus, *Puccinia striiformis* f. sp. *tritici* (*Pst*), is an economically important disease that affects global wheat production. This disease manifests on wheat leaves and leads to severe yield losses, potentially amounting to a billion dollars a year globally (estimated in 2015 to $979M per year) (Beddow et al. 2015) when not controlled with fungicide applications (Chen 2014). Deploying genetic resistance is preferable for efficiency and reduced environmental impact, but requires identification of new resistance alleles to combat pathogen evolution. Over the last decade, the severity of the disease has increased due to the emergence of new virulent races in East Asia and Africa (Hubbard et al. 2015), the centers of wheat origin and genetic diversity (Vavilov and Dorofeev 1992). In other areas, the dramatic shift of the pathogen population, including introduction of the new *Pst* ‘Warrior’ race in the United Kingdom, has triggered widespread epidemics (Hubbard et al. 2015). Newer isolates of *Pst* are now found in wider geographic locations and have become more adapted to warmer and drier climates. Climate change is expected to facilitate the spread of *Pst* and exacerbate the vulnerability of wheat. New sources of disease resistance are urgently needed as current varieties fall behind against newly emerging pathogen races. Over the last 50 years, the geographic range and severity of stripe rust has increased such that 88% of the world’s wheat is susceptible to rust. Since 1.2 billion people depend upon wheat for their calories, it has been estimated that $32M USD is a well-justified annual expenditure in the development and deployment of stripe rust-resistant varieties (Beddow et al. 2015).

Yellow rust (*Yr*) resistance genes have been central in developing varieties with enhanced disease resistance and are widespread in commercial varieties (Marchal et al. 2018). Nevertheless, phenotypic expression of stripe rust resistance varies substantially as evidenced by the eight cloned *Yr* genes (Yuan et al. 2018; Singh et al. 2023). Intracellular immune receptor *Yr5* (BED-NLR) gives complete resistance with no visible infection (Wang and Chen 2017), ABC transporter *Yr18* plants frequently exhibit leaf tip necrosis (Shah et al. 2011), and plants carrying hexose transporter *Yr46* often have leaves striped with chlorotic and necrotic tissue (Moore et al. 2015). The expression of these phenotypes is further influenced by genetic background, the genotype of the *Pst* isolate and environmental conditions (Wang and Chen 2017). Currently, rampant spread of *Pst* isolates of the highly virulent molecular group five (MG5), present in the United States, can only be restricted by the ubiquitously used broad-spectrum stripe rust-resistance genes *Yr*5 and *Yr15*, plus a few newly identified *Yr* genes. Therefore, the need to release new resistant varieties, especially those with durable, non-race specific resistance, is urgent before new races emerge to overcome these genes (Sharma-Poudyal et al. 2020). A race (TSA-6) was reported as virulent to *Yr5* in China in 2017 (Zhang et al. 2022b). Similarly virulent *Pst* isolates to *Yr5* have been reported in India, China, Australia, and Turkey (Tekin et al. 2021).

The diversity in protein function of the *Yr* genes are indicative of the multiple pathways through which resistance can occur. Development of necrotic lesions through activating a hypersensitive immune response at the point of pathogen infection is typical of race-specific resistance (Balint-Kurti 2019). Mutations affecting genes in the programed cell death pathway can create lesions that form without pathogen exposure, leading to the ‘lesion mimic’ phenotype, long known in maize (Hoisington et al. 1982) and other grasses (McGrann et al. 2015). Cloned genes that give these phenotypes include several alleles of the *Rp1* (resistance to *Puccinia sorghi*) locus (Hu et al. 1996; Collins et al. 1999), of these *Rp1-D21* forms spontaneous lesions suppressed by temperatures above 30°C (Negeri et al. 2013). A similar suppression of lesion phenotype has been described in the rice ethyl methanesulfonate (EMS) mutant, *white and lesion mimic leaf1* which is expressive at 20°C but suppressed at 28°C (Chen et al. 2018a). Another mechanism by which autoimmune leaf lesions can form is by combining NLR (Nucleotide Binding Leucine Rich Repeat) resistance (R) proteins with non-coadapted host proteins. An example of this is spontaneous lesion formation in interspecific hybrid (*Solanum lycopersicum* x *Solanum pimpinellifolium*) tomato leaves (Rooney et al. 2005). EMS mutagenesis at multiple antagonistic loci could spawn lesion-mimic phenotypes by creating similarly maladapted immunity proteins.

*Pst* often co-occurs with other major wheat fungal pathogens, some of which can have opposite lifestyles requiring careful selection of breeding material for responsible deployment. Stripe rust and blotch diseases have overlapping geographical ranges and are both favored by cool temperatures (Smith et al. 2021). Thus, we evaluated the EDR lines for their response to both biotrophic and/or necrotrophic fungal pathogens at the seedling and adult plant stages. Moderately resistant wheat genotypes select for pathogens having increased aggressiveness, suggesting the need for varieties with combined resistance to both blotch and rust (Cowger and Mundt 2002).

Wheat has a large polyploid genome which provides both unique advantages and challenges in genetic identification of new beneficial alleles in forward screens. Polyploidy results in multiple gene copies. The resulting gene redundancy buffers the effects of deleterious alleles allowing for a higher density of mutations without severely affecting plant health but, it also complicates the analysis of the effects of individual single mutations. Over 2,000 lines from an EMS-mutagenized tetraploid elite wheat *Triticum turgidum* subsp. *durum* cv ‘Kronos’ population have been characterized by exome capture and sequencing (Uauy et al. 2009; Krasileva et al. 2017). The mutations in this population have been useful toward many breeding (Mo et al. 2018; Harrington et al. 2019; Li et al. 2021; Zhang et al. 2023a) and basic research goals (Schilling et al. 2020; Wang et al. 2020b; Corredor-Moreno et al. 2022). Mutations in the population are incorporated into multiple plant genome databases and mutant seed is available from several seed banks (Krasileva et al. 2017). We leveraged these unique resources to identify previously undescribed germplasm with enhanced fungal disease resistance.

In this paper, we identified 16 lines that showed persistent enhanced adult plant stripe rust resistance over four years of field trials. In addition, we documented their seedling responses to *Pst*. To assess additional gain of resistance, we evaluated these EDR lines for their response to other wheat pathogens including *Puccinia triticina* causing leaf rust, *Blumeria graminis* f. sp. *tritici* causing powdery mildew and *Pyrenophora tritici-repentis* causing tan spot at the seedling stage as well as to *Zymoseptoria tritici* causing Septoria tritici blotch (STB) and *Fusarium graminearum* causing fusarium head blight (FHB) at the adult plant stage. Five of the persistently resistant lines were determined to have autoimmune and cold-enhanced lesion mimic phenotypes. EMS mutations previously sequenced with exome capture were mapped to a new long-read Kronos genome assembly to increase the utility of these lines in breeding and genetic studies.

## Results

### Sixteen EDR lines are persistently resistant to stripe rust

A total of 2,000 independent Kronos EMS-mutagenized M3 families (Uauy et al. 2009; Krasileva et al. 2017) were screened for resistance to stripe rust in spring growing seasons in Davis, California, with half of the mutants grown from 2012 to 2013 and the other half from 2013 to 2014. Field-collected *Pst* urediniospores from the previous season were used to inoculate highly susceptible border plants (Fig. 1a) and provide high pathogen pressure for effective plant resistance screening. M4 heads were collected from these lines and bulked for seed for future screening. M5, M7, M8 and M9 lines were grown again and scored for stripe rust resistance (Table S1) over multiple years. Wild type Kronos has intermediate stripe rust resistance at the adult plant stage. Based on 0-9 infection type (IT) scale (McNeal FH 1971), ITs of 0-3 indicates that the pathogen has not formed uredinia, wild type Kronos had an average IT of 4 (ranging from 3 to 5) over both screening years with visible stripe rust uredinia appearing in necrotic stripes occupying ∼10% of the flag leaf area. We chose a threshold of having an average IT of less than 3 in 2014 and 2019 to 2021 (and ranging from 1 to 4) (Table S1) to categorize the lines as EDR lines having ‘persistent resistance.’ Of the 34 candidate EDR mutant lines originally identified as resistant in years 1 and 2 of the screen (Table S2), 16 lines met the threshold of persistently resistant and of these five had autoimmune lesion mimic phenotypes (Fig. 1).

**Figure 1.**
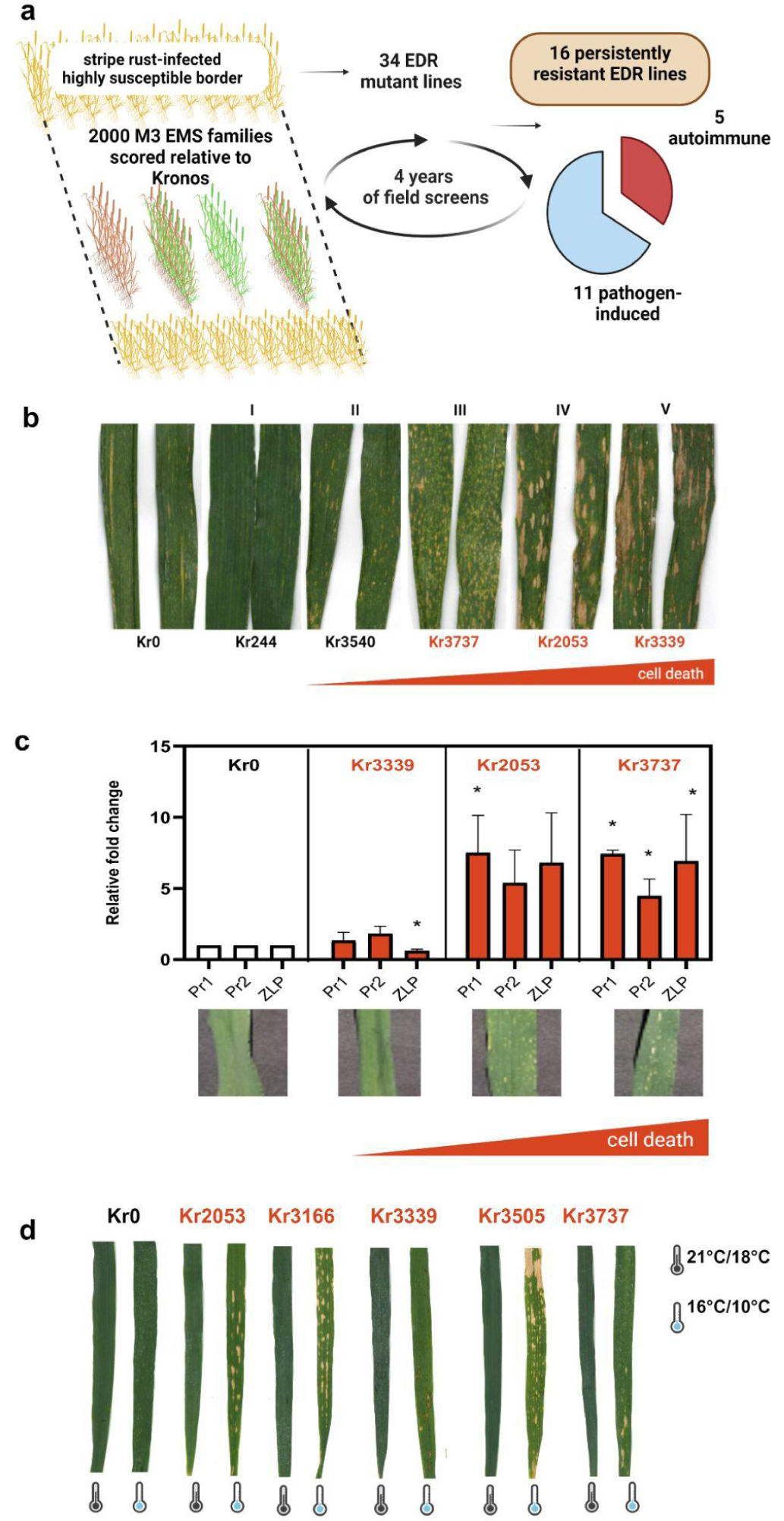
Diverse Kronos EDR stripe rust mutants. a) Identification of stripe rust EDR lines. Over two growing seasons, 2000 EMS M3 lines were planted in 30 cm rows with border rows of highly susceptible genotypes and were selected on their ability to prevent pathogen sporulation, unlike the wild-type progenitor Kronos. b) Excised flag leaves of Kronos (left) and phenotypic classes with increasing levels of macroscopic cell death in rust field-grown mutants fit into 5 observed phenotypic categories (I-V). Kr244, a non-autoimmune “green” EDR mutant is included for comparison (left) to representative mutants Kr3540, Kr3737, Kr2053 and Kr3339. c) Relative fold change of rust-inducible genes Pr1, Pr2 and Zlp correlates with autoactive lesion formation in LMM mutants. d) Autoactive lesioned EDR mutants display cool temperature-enhanced cell death in the absence of the stripe rust pathogen. About 10% of putative EDR lines grown in fungicide-treated field plots in 2014 formed lesions beginning at GS 37-39 after sowing (Zadok 1974). This phenotype was confirmed to be spontaneous and enhanced by cool temperatures, by growing the lines under normal greenhouse conditions (21°C/18°C day/night (+/- 2°C) 16/8 hr) and in a cool growth chamber (16°C day/10°C night; 16hr daylength).

Pathogen-challenged flag leaves of EDR mutant lines showed a spectrum of response from no macroscopic cell death lesions to lesions covering more than half of the leaf (Fig. 1b; Fig. S1). Kronos has moderate adult plant susceptibility (APS) and forms necrotic lesions with a light amount of rust sporulation. One line, Kr244, had no macroscopic cell death, but others showed large coalescing lesions affecting most of the leaf area. In our rust nursery in Davis, CA, USA the 16 persistently resistant mutants fell into five major phenotypic classes; I: no macroscopic cell death (n=1; Kr244), II: moderate cell death covering up to 30% of the leaf surface (n=13; Kr3540 is shown), III: extensive cell death with many tiny lesions speckling 80% or more of the leaf (n=1; Kr3737), IV: extensive cell death due to large oval lesions at the leaf tip and medium lesions in the proximal half of the leaf (n=1; Kr3540) and V: extensive cell death due to large, striped coalescing lesions beginning in the leaf center (n=1; Kr3339) (Fig. 1b; Fig. S1). Within the EDR lines, only Kr244 lacked lesions, so whether they developed as a consequence of pathogen exposure or spontaneously, they were associated with enhanced resistance to stripe rust. Most of the variation in lesion pattern and size is within the autoimmune lines. The EDR lines that form lesions only in the presence of the pathogen all fall into phenotypic category II. Early descriptions of *Les* mutants in maize (Hoisington et al. 1982) described established lesions as having two components: initiation and enlargement. Kr3737 (category III) was high in lesion initiation, having many pinprick-sized lesions that gave the leaves a speckled appearance, but low in lesion expansion. Whereas Kr3339 (V) had fewer but quite expansive lesions, Kr2053 (IV) was intermediate in its expression of both components.

To explore the relationship between lesion formation and stripe rust resistance in the autoactive lines, we compared expression levels of 3 genes, Pr1-1-1 (*TraesCS5B02G181500*), Pr-2-1c (*TraesCS3A02G483000*), and a Zeamatin-like protein (Zlp) gene (*TraesCS4A02G29630*) that were identified as being *Pst*-inducible ((Zhang et al. 2014b) and Bioproject PRJNA387101 at NCBI (https://www.ncbi.nlm.nih.gov/bioproject/PRJNA387101)). Levels of the pathogenesis related gene expression roughly correlated with the severity of lesion formation (Fig. 1c), indicating an overlap in the induction of these processes.

The original screen was conducted in Davis, CA in *Pst*-inoculated plots without fungicide treatment, which allowed us to observe pathogen-dependent phenotypes at relatively high temperatures (the average April daily temperature in Davis in 2013-2014 was 17°C with an average low of 9°C and an average high of 24.9°C (www.weather.gov)). The Kronos mutant population was grown and observed again in June of 2015, in Norwich, UK, in a fungicide-treated plot to increase seed. By contrast, the temperature averages in Norwich over the observation period were cooler (the average June daily temperature in Norwich, UK in 2015 was 14.2°C with an average low of 8.6°C and an average high of 19.8°C). From the original candidate list of 34 resistant lines (Table S2), 9 autoimmune M4 lines were identified as having lesions or chlorosis under the cooler conditions in Norwich. For confirmation of these phenotypes, and assuming that these phenotypes would be enhanced in daytime temperatures below 20°C, as seen in similar mutants (Fu et al. 2009; Negeri et al. 2013; Chen et al. 2018a), the candidate autoimmune lines and Kronos were grown in growth chambers under cool conditions (16°C/10°C day/night 16/8 hr) and greenhouse conditions (21°C/18°C day/night (+/- 2°C) 16/8 hr) without rust pressure, and the sensitivity of the phenotype to cool temperatures was confirmed. Under standard conditions, of the persistently resistant 16 lines, distinct lesions formed in 5 lines (Kr2053, Kr3166, Kr3505, Kr3737, and Kr3339) (Fig. 1d; Fig. S1a), and 8 of the 9 originally identified lines had more pronounced visible lesions under cooler temperatures. Kr456 did not form lesions under either tested condition (Fig. S1b).

Sixteen EDR lines were highly resistant to stripe rust by fitting our threshold of having an average IT of <3 and preventing pathogen sporulation in the adult-plant stage in rust-inoculated fields over four years. Of these 16 lines, five lines with an IT of no more than 2 and seven lines with an IT no more than 3 were significantly more resistant (*p* < 0.01, and *p* < 0.05, respectively) to stripe rust than Kronos at the adult stage, and the other four lines with an IT of no more than 4 were not significant but had an IT lower than Kronos (Fig. 2a) by ordinary one-way ANOVA, (*p* < 0.05). Since the original disease screen of EMS mutants was conducted on adult plants, most EDR lines were similarly susceptible as Kronos at the seedling stage. An exception was Kr3186 which showed high seedling resistance to race PSTv-37 in greenhouse trials (Fig. 2b). All the other EDR lines that differed from Kronos had lower average IT scores than Kronos (Fig. 2a). We have isolated EDR EMS lines with lower IT than wild type Kronos, suggesting that we can achieve resistance through modulating several distinct genes and pathways. Hussain et al. 2018, created a hexaploid wheat EMS population from cultivar NN-Gandgum-1, also moderately resistant to stripe rust and, found twenty EDR lines from 3624 M2 plants with an efficiency of 0.55%. We obtained 0.8% efficiency in a Kronos mutagenized population, indicating that forward screens for stripe rust resistance in EMS wheat populations is an effective strategy.

**Figure 2.**
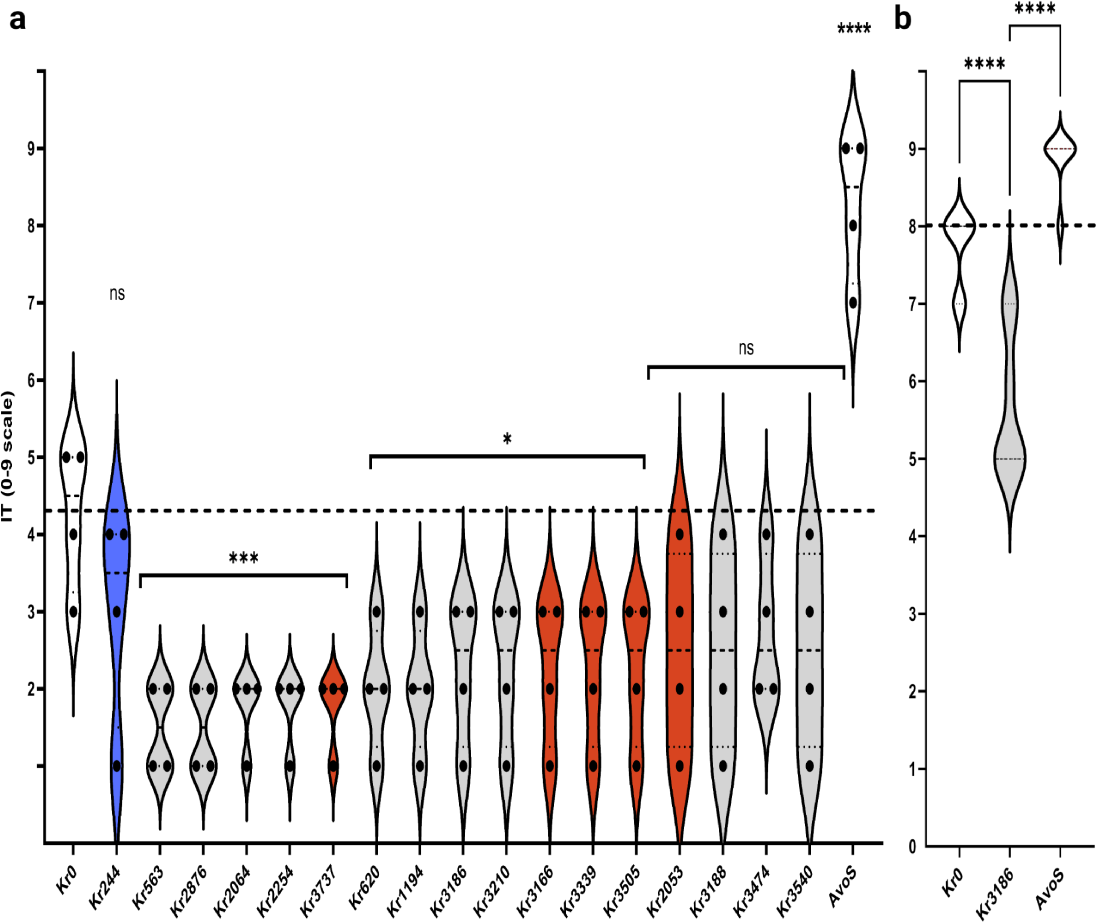
Diverse Kronos EDR stripe rust mutants are persistently resistant. a) EDR Adult stripe rust IT. Sixteen EDR lines are more resistant than the progenitor variety, Kronos to Davis, CA rust races over four years at the adult stage. Each dot represents a year (2014, 2019-2021, in Davis, CA, USA). All presented lines had an average score of less than 3 and prevented pathogen sporulation. b) EDR seedling stripe rust IT against stripe rust race PSTv-37. EDR line Kr3186 had the lowest stripe rust IT score. AvoS was the susceptible check variety in both experiments. Infection types (IT) of all plants are scored on the 0-9 scale (McNeal, 1971). Control lines (white) plus an EDR line with no macroscopic cell death under pathogen challenge (blue) are included for comparison with the autoactive lesioned lines (red). Dotted line indicates Kronos median value of 4.25 in the adult stage experiment and 7.75 in the seedling stage experiment. Means were compared using one-way ANOVA, *, significant at *p* = 0.05; **, significant at *p* = 0.01; *** significant at *p* = 0.001; ****, significant at *p* < 0.0001; ns, non-significant.

### Autoimmune EDR stripe rust mutants showed an all-stage trade-off to foliar necrotrophic pathogens but not to FHB

To explore if the EDR lines have differential adult responses to other pathogens, the 16 lines were grown in a field highly infected with the *Septoria tritici* blotch (STB) pathogen, *Z. tritici,* in Corvallis, Oregon in replicated trials in 2021 (M8) and 2022 (M9). The EDR lines were sown in October to expose them to fall ascospore infections and subsequent infections via conidia in winter and spring. Spring wheat genotypes, such as Kronos, typically survive the mild winters of Corvallis. The lines were compared with the locally adapted winter cultivars Bobtail, Madsen, and Stephens, which are moderately resistant, moderately susceptible, and susceptible to STB, respectively). Scoring for IT was on a subjective 1 (little or no disease) to 5 (highly diseased) scale (Fig. S3b and Fig. S4). Kr244, which has little or no macroscopic cell death and is not autoimmune (Fig. 1b), performed better than Kronos in the STB nursery as shown by their average ranked scores over both years (Fig. 3a). Kronos showed characteristic pycnidial formation (Fig. 3a, white and blue arrow). Four lines performed worse than Kronos and two of these have an autoimmune lesioned phenotype that is enhanced by cool temperatures (Fig. 1d and Fig. S1), which are typical of the early Oregon wheat growing season (during October - June). These lines have leaves with necrotic tissue patches, favored by the STB pathogen. All of the mean STB scores for the autoimmune lines were higher than that of Kronos (Fig. 3a; Table S2).

**Figure 3.**
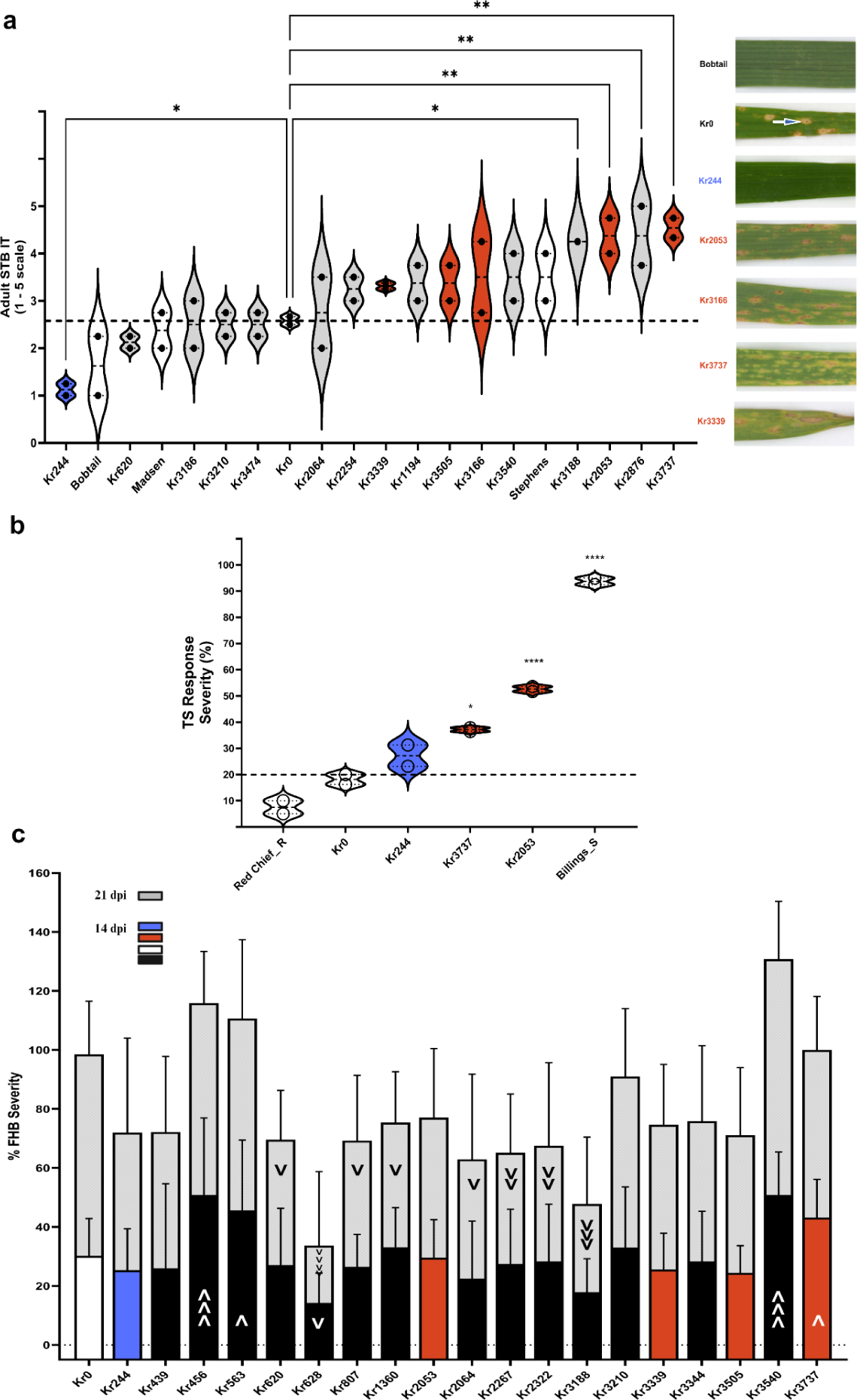
Autoactive lesion mimic EDR mutants are less resistant to foliar necrotrophic pathogens. a) (Left) Violin plot of ranked STB response (1-5 scale) of EDR lines shows trend that autoimmune lesioned lines have higher IT scores than non-autoimmune lines. Each dot represents a year. (Right) Excised flag leaves collected in STB nursery. Bobtail (resistant check), Kronos with STB diagnostic black pycnidia (white arrow), Kr244 (blue) and four autoimmune lesion mimic plants (red). b) Ranked seedling tan spot severity shows a trend that autoimmune lesion mimic EDR mutants have higher severity. (A-B) Dotted line indicates Kronos median value. Means were compared using one-way ANOVA, *, significant at *p* = 0.05; **, significant at *p* = 0.01; *** significant at *p* = 0.001; ****, significant at *p* < 0.0001; ns, non-significant. c) FHB severity (%) at 14 dpi (lower bars) and 21 dpi (upper gray bars) are shown. Significant differences are shown as “v” for less than or “upside-down v” for more than Kronos. Having one symbol indicates significance by one-way ANOVA at the *p* = 0.05 level, two symbols indicate significance at the *p* = 0.01 level and three symbols indicate significance at the *p* = 0.001 level. Control lines (white) plus an EDR line with no macroscopic cell death under pathogen challenge (blue) are included for comparison with the autoactive lesioned lines (red) and the non-autoimmune lines (black).

To further investigate a possible sensitivity of the autoimmune lines to necrotrophic pathogens, we assayed their seedling response to the tan spot pathogen, *P. tritici-repentis*. The autoimmune lines Kr3737 and Kr2053 had higher severity (%) than Kronos or Kr244 (Fig. 3b), even before the lesion phenotype was expressed. This suggested that even at the seedling stage, the lesions may confer enhanced susceptibility to tan spot. We saw a similar result when we challenged these lines with biotrophic *P. triticina* races MNPSD and MPPSD (Fig. S3b). When seedlings were challenged with *P. striiformis* f. sp. *tritici* PSTv-37, *P. triticina* race BBBQD, and *B. graminis* f. sp. *tritici,* Kr3737 and Kr2053 were not more sensitive than Kronos (Fig. S2b; Fig. S2c, respectively) which showed large variation in response. A possible explanation is that there are interactions between alleles in the individual lines and the different pathogens.

We also assayed the EDR lines for response to another adult stage-infecting hemibiotrophic/ necrotrophic pathogen, *F. graminearum,* which causes FHB of wheat, in the greenhouse experiments. The 16 persistently stripe rust resistant EDR lines showed differential responses to *F. graminearum* inoculation both at 14 dpi (days post-inoculation) and at 21 dpi. Unlike their responses to the foliar diseases (STB, leaf rust and tan spot) which tended to be worse in mutants with autoimmune leaf necrosis (Fig. 3a, Fig. S2b, Fig. S4), three of the four autoimmune mutants tested were not significantly different from Kronos (Fig. 3c). At 14 dpi, the average FHB severity for susceptible Kronos was 30.2%. At 21 dpi, Kronos exhibited an average severity of 68.3%. Kr3188 showed high resistance at 21 dpi and was not significantly different from Cadenza (one-way ANOVA, *p* = 0.5581). At the 21 dpi time point, two lines, Kr2267 and Kr2322, had significantly lower severity at p<0.01 and four lines (Kr628, Kr2267, Kr2322, and Kr3188) had significantly lower severity at *p* < 0.05. Four EDR lines, Kr456, Kr563, Kr3540 and Kr3737 had significantly higher severity, all at 14 dpi (Fig. 3c). Altogether, our results showed that autoimmune lines tended to fare worse when exposed to STB but that even though the EDR lines varied in their response to FHB, taken together, there was no strong positive or negative correlation between stripe rust resistance and FHB resistance.

### Autoactive lesioned EDR mutants do not show reduced TKW

To assess possible grain mass penalties due to autoimmunity or other mechanisms of enhanced resistance, we measured thousand kernel weights (TKW) in the EDR lines in our stripe rust trials. Most EDR lines had indistinguishable TKW values from Kronos (Fig. S5), except for three lines that had TKWs significantly higher than Kronos (one-way ANOVA, *p* = 0.05). Two lines Kr2053 and Kr3339, formed lesions with and without the rust pathogen, suggesting that in the presence of this pathogen, these lines are not inherently at disadvantage due to defense priming nor ectopic, localized cell death lesions. More yield measurements would be needed in the future using recombinant inbred lines isogenic for the LMM mutations to test this.

### Re-mapping of EDR line exome data to new Kronos long-read assembly

A chromosome-level assembly of the Kronos genome has recently become available (Seong et al. 2023 https://zenodo.org/records/11106422). In support of forthcoming functional research on EDR mutants, we remapped their exome capture data onto this reference genome and identified single nucleotide polymorphisms (SNPs) with the MAPS pipeline (Table S9a) (Henry et al. 2014; Krasileva et al. 2017). We excluded two lines, Kr2027 and Kr2067, from our analysis due data unavailability.

To discern high-confidence mutations, our initial criteria required a minimum coverage of five mutant reads for heterozygous (HetMC) mutations and three for homozygous (HomMC) mutations, adhering to the methodology in the previous study (Krasileva et al. 2017). At this stringency, the MAPS pipeline identified only two substitutions from a non-mutagnized control line (Kr0), indicating no essential variation between the reference genome and the control line (Fig. 4a). Analysis of 32 EDR lines revealed 89,120 substitutions with 88,343 (99.1%) characterized as EMS-type transitions (Table S9a). These numbers surpass those reported using the hexaploid Chinese Spring genome as a reference (Krasileva et al. 2017). To maximize the number of detected mutations while maintaining the targeted EMS-type mutation rate of 98%, we adjusted the stringency of HomMC and HetMC as in the most recent study (Zhang et al. 2023b). We could identify 14,692 additional EMS-type mutations from the 32 EDR lines, averaging 3,220 total EMS-type mutations per line (Fig. 4a and Fig. S6; Table S9b).

**Figure 4.**
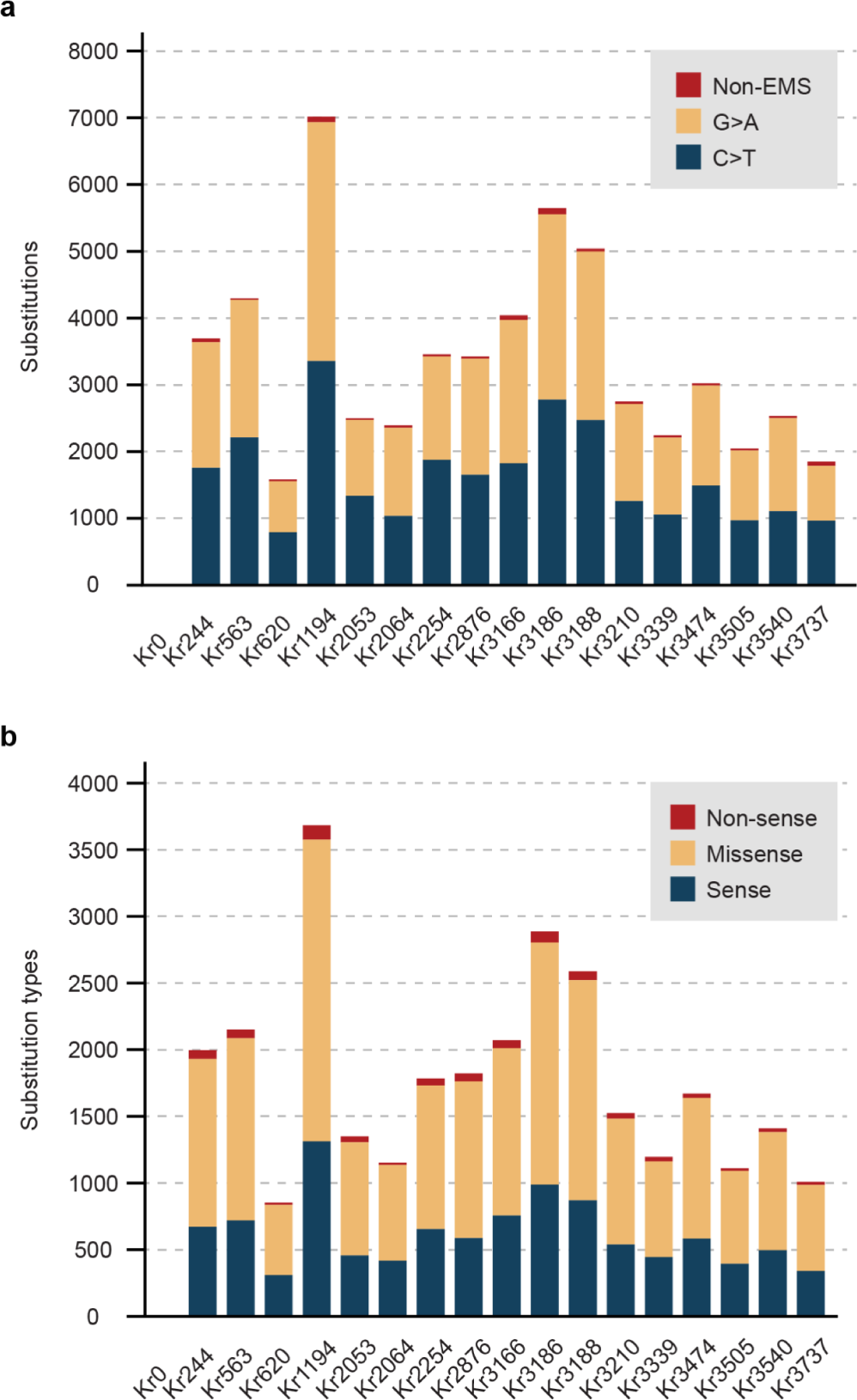
Identified substitutions in EDR lines and their impacts on proteins. a) The number of substitutions identified with the MAPS pipeline from major EDR lines and non-mutagenized control line Kr0. The mutations are colored to indicate non-EMS type mutations, and two EMS-type mutations (G to A and C to T). b) The mutation effects of identified substitutions in coding sequences.

We examined effects of the identified mutations on each EDR line, using 69,808 high-confidence genes (Seong et al. 2023). For each gene, we chose the annotated transcript producing the longest protein product. On average, 77% of all identified mutations impacted annotated genic regions, including coding sequences, splicing regions, introns and untranslated regions (Fig. S7). Approximately 7% of mutations were located in potential regulatory regions near genes, while 16% in intergenic regions. On average, mutations affected 1,654 genes per line, introducing 596 synonymous, 1,065 missense, and 46 nonsense mutations in coding sequences (Fig. 4b; Table S9c).

We investigated mutations within orthologs of known lesion mimic mutants (LMMs), suppressors of LMMs, and *Yr* gene products (Table S10a) (Fu et al. 2009; Krattinger et al. 2009; Moore et al. 2015; Marchal et al. 2018; Zhang et al. 2019; Wang et al. 2020a; Klymiuk et al. 2020). Across the 32 EDR mutant lines, we identified 221 missense mutations and 5 nonsense mutations in 134 orthologs. Some of these mutations may influence the phenotypes observed in the EDR mutants (Table S10b). In the five lines—Kr2053, Kr3166, Kr3505, Kr3737 and Kr3339—which exhibited distinct lesions in the absence of pathogens, we detected 22 missense mutations in 21 orthologs of LMMs. Kr2053 and Kr3339 both have missense mutations in the Kronos ortholog of *Arabidopsis* candidate gene nonsense-mediated decay factor UPF1 (Table 1). This comprehensive mapping of mutations in recognized EDR genes from the Kronos genome will facilitate future reverse genetic research in these lines.

**Table 1.** Substitutions within coding sequences of orthologs of LMMs. Mutant genes inducing lesion mimic phenotypes were identified in *Arabidopsis thaliana* and *Oryza sativa*. Their orthologs in Kronos were investigated in the five EDR lines exhibiting autoactive lesion phenotypes and listed if missense mutations are present. No nonsense mutations were identified in the surveyed orthologs of all lesion mimic mutant genes.

| Mutant line | Proteins in Kronos |  |  |  | Putative ortholog in Chinese Spring | Reference |  |
| --- | --- | --- | --- | --- | --- | --- | --- |
|  | ID | Position | Reference | Mutated |  | ID | Common name |
| Kr2053 | TrturKRN1A01G038710.1 | 397 | R | K | TraesCS1A02G259600.1 | AT5G48380 | BIR1 |
| Kr2053 | TrturKRN1A01G039980.1 | 118 | P | S | TraesCS1A02G270000.1 | Os10g0195600 | SPL18 |
| Kr2053 | TrturKRN2A01G034020.4 | 1170 | P | S | TraesCS2A02G228000.1 | AT5G47010 | UPF1 |
| Kr2053 | TrturKRN2B01G044330.1 | 961 | G | S | TraesCS2B02G258400.1 | Os07g0203700 | SPL5 |
| Kr2053 | TrturKRN6A01G001770.2 | 200 | A | V | TraesCS6A02G006400.1 | AT1G55490 | LEN1 |
| Kr2053 | TrturKRN7B01G031430.2 | 274 | G | W | TraesCS7B02G201000.1 | AT4G01370 | MAPK4 |
| Kr3166 | TrturKRN2A01G062630.1 | 145 | P | S | TraesCS2A02G437600.1 | Os04g0601800 | WSP1 |
| Kr3166 | TrturKRN4B01G030810.1 | 13 | P | L | TraesCS4B02G188200.1 | AT4G33680 | AGD2 |
| Kr3166 | TrturKRN6A01G004900.1 | 416 | G | D | TraesCS6A02G024300.1 | Os03g0364400 | BBS1 |
| Kr3166 | TrturKRN6A01G038280.1 | 383 | G | E | TraesCS6A02G244000.1 | AT5G51290 | ACD5 |
| Kr3166 | TrturKRN7A01G005700.5 | 343 | G | C | TraesCS7A02G036200.2 | AT1G55490 | LEN1 |
| Kr3339 | TrturKRN1B01G046720.1 | 70 | D | N | TraesCS1B02G269900.1 | AT5G48380 | BIR1 |
| Kr3339 | TrturKRN2A01G033500.2 | 514 | T | I | TraesCS2A02G231900.3 | AT1G02120 | VAD1 |
| Kr3339 | TrturKRN2A01G034020.4 | 584 | E | K | TraesCS2A02G228000.1 | AT5G47010 | UPF1 |
| Kr3339 | TrturKRN5B01G027370.1 | 818 | P | L | TraesCS5B02G160100.1 | AT5G19400 | SMG7 |
| Kr3339 | TrturKRN6B01G057640.2 | 408 | G | D | TraesCS6B02G358000.3 | AT5G64930 | CPR5 |
| Kr3505 | TrturKRN1A01G012390.2 | 358 | V | I | TraesCS1A02G065700.2 | AT5G60410 | SIZ1 |
| Kr3505 | TrturKRN1B01G052190.1 | 335 | A | T | TraesCS1B02G307200.1.cds1 | AT5G58430 | EXO70B1 |
| Kr3505 | TrturKRN3A01G047940.1 | 558 | G | D | TraesCS3A02G348200.1.cds1 | AT5G58430 | EXO70B1 |
| Kr3505 | TrturKRN5A01G024990.1 | 53 | P | S | TraesCS5A02G175200.1 | AT3G14110 | FLU |
| Kr3505 | TrturKRN5A01G029260.1 | 147 | S | F | TraesCS5A02G205900.1 | AT3G25540 | LOH1 |
| Kr3737 | TrturKRN2B01G073290.1 | 653 | A | T | TraesCS2B02G465300.1 | Os01g0116600 | SPL33 |

## Materials & Methods

### Field trials for adult fungal infection response

#### Identification of stripe rust EDR lines and evaluation over multiple growing seasons

The initial forward genetic screening of an EMS mutant population of the wheat variety Kronos for *Pst* resistance was conducted in the fields at the UC Davis Field Station in Davis, California during spring of 2013 and 2014. In each of these two seasons 1,000 independent M3 families were evaluated. In all field assessments, each family or genotype was planted in a single 1 m long row, spaced 30 cm apart. To ensure sufficient inoculation pressure, these plots were surrounded by a *Pst* susceptible breeding line, D6301. At plant growth stage GS 40-49 (booting) (Zadoks et al. 1974), the plots were inoculated with *Pst* urediniospores collected from the previous field season. *Pst* infection type (IT) was scored on a 0-9 scale (McNeal FH 1971) 2-4 times per season, separated by 7-10 days. In both years, non-mutagenized Kronos had an IT of 4 to 5. This intermediate resistance allowed for the identification of mutant families that displayed enhanced resistance. After sporulation was observed on the flag leaf of wild-type Kronos, the entire population was observed at least twice for families that had plants that deviated from the Kronos IT, mutants that did not display any visible sporulation were identified as candidate EDR lines. Single tillers of plants displaying deviant phenotypes were labeled and the corresponding spike was harvested at maturity for subsequent evaluation. For evaluation over multiple growing seasons, seeds of the EDR lines were planted in 1 m long rows, surrounded by a highly susceptible variety and inoculated with *Pst* spores collected from the previous years. The EDR lines and Kronos were grown with identical spacing as in their original screening trials, but in triplicate randomized blocks.

#### Chamber Validation of stripe rust EDR phenotypes

EDR lines with strong resistance phenotypes were included in validation experiments in controlled environment growth chambers and inoculated with *Pst*. To ensure a fresh and robust inoculation pressure, the highly *Pst* susceptible lines Avocet S and Fielder were grown and inoculated with spores collected from the field in advance of planting the experiment. The experimental plants were inoculated at mid-tillering phase and again after flag leaf emergence. Freshly collected spores were mixed with technical grade talcum powder and applied to plants that had been misted with deionized water and kept in a dark dew chamber at 10°C for 24 hr. The temperature conditions of the chamber were then set to gradually increase from a nighttime temperature of 4°C to 18°C during the days with a photoperiod of 16 hr. Pustules of *Pst* began to appear 14 days after inoculation and phenotypes were recorded 25 to 30 days after the inoculation.

### High-throughput assessment of thousand kernel weight (TKW) under stripe rust exposure

For TKW assessment, 200-300 kernels of each EDR line and Kronos of each of three biological replicates were averaged over the three years, 2020 to 2022 in Davis, CA. Kernels were distributed evenly in a (11.3 cm x 13.1 cm) box, weighed and photographed with a 9th generation iPad camera (model A2197). The number of seeds in each photograph was determined using the CountSeeds App with the Wheat Seed template and the average weight per kernel was then determined. This number was multiplied by 1000 to give the estimated TKW. Three technical replications were made of each biological replicate, without replacement.

### Identification and validation of autoimmune EDR mutants

Nine autoimmune mutant lines were originally identified as having lesions in a fungicide-treated plot (rust-free) in Norwich, UK, 2015. For confirmation of these phenotypes, this subset of lines and Kronos was grown in 3 inch pots using Supersoil #4 supplemented with Osmocote 14-14-14 at 3.3g/L and grown in a growth chamber under cool conditions (16°C/10°C day/night 16hr/8hr). Eight of the nine identified lines had lesions under these conditions, but none had lesions under standard greenhouse conditions (21°C/18°C day/night (+/- 2°C) 16/8 hr).

### Real-time quantitative PCR (RT-qPCR) analysis

Kronos, Kr2053, Kr3339, and Kr3737 were grown in 8” plastic pots (1.4L) in LM-6 soil (Lambert, Canada) in the greenhouse at 22°C/16°C day/night with a 16 hr photoperiod. Flag leaves (Zadok’s GS 55) were collected in three replicates from each accession and immediately frozen in liquid nitrogen. Total RNA was extracted using TRIzol reagent (Invitrogen, cat. no. 15596018) according to the manufacturer’s instructions. DNA contamination was removed by treating total RNA samples with DNase I (Roche, Cat. No. 04716728001), followed by phenol-chloroform purification and precipitation. First-strand cDNA was synthesized using the SuperScript™ IV first-strand synthesis kit (Invitrogen, Cat. No. 18091200). RT-qPCR was then performed in four technical replicates for each sample with the PowerUp™ SYBR™ green master mix (Applied Biosystems, Cat. No. A25741) on the QuantStudio™ 5 Real-Time PCR System, 384-well (Applied Biosystems, Cat. No. A28140) with the following program: UDG activation at 50℃ for 2 min, activation (Dual-Lock™ DNA polymerase) at 95℃ for 2 min, followed by 40 cycles of denaturing at 95℃ for 15 sec, annealing/extension at 60℃ for one min. Relative expression levels of Pr1, PR2 and ZLP were calculated using the comparative cycle threshold (2^−ΔΔCt^) method (Livak and Schmittgen 2001) with normalization to the internal control gene *Actin*. Raw data and calculations are provided in Table S3. All primers used for RT-qPCR are shown in Table S8.

### Adult plant screens toward necrotrophic pathogen response

#### Field trials for STB response

In our 2020 pre-trial of Kronos and a smaller set of tester EDR lines, we had aphid-transmitted barley yellow dwarf virus pressure. Hence, in 2021 and 2022 we applied Cruiser Maxx Vibrance Cereals (Syngenta) at a rate of 0.48 mL per g of seed and included all available lines and controls. The EDR lines were grown in four replicates at the naturally infected STB plot in Hyslop Field in Corvallis, OR, in a randomized complete block design. STB infection type was scored on a subjective 1 (resistant) to 5 (susceptible) scale using moderately resistant, moderately susceptible, and susceptible check varieties Bobtail, Madsen and Stephens, respectively.

#### Greenhouse tests of FHB severity

Kronos wild type was used as a control and EDR lines were tested using five plants per genotype planted in a randomized complete block design in 4 inch x 4 inch pots in the greenhouse, in the following soil mix: Sungro professional mix SS#1, Metromix 830 and Surface Greens grade (2:2:1). Temperature conditions were 23-25°C during daytime and 16-18°C at night with 16 hr of light. Highly virulent *F. graminearum* isolate GZ3639 (Desjardins et al. 1997) was used as the inoculum. To generate macroconidia, 2-3 fungal mycelial culture plugs from Potato Dextrose Agar were introduced into Mung bean broth (MBB). Macroconidia were counted using a hemocytometer, and an inoculum was created by diluting the culture to 1 × 105 spores/ml using sterile water. Inoculations were conducted at the pre-anthesis stage, approximately two days before anthers emerged from the spikes. The 10th and 11th spikelets, counted from the base of the spikes, were marked. Ten microliters of inoculum were pipetted into one floret of each spikelet between the lemma and the palea, taking care to avoid injury to the other parts of the florets. Following this, the spikes were enveloped in moisture-saturated plastic bags for 72 hr to create a high-humidity environment conducive to optimal fungal growth (Chhabra et al. 2021b). Bleached spikelets were counted from the point of inoculation downward, and % severity was calculated by dividing bleached spikelets by total number of spikelets beneath the inoculated spikelet (10) at 14 dpi and 21 dpi.

### Seedling screens to multiple pathogens

#### Stripe rust

Five to seven plants of each EDR line or Kronos, and the susceptible check Avocet S were grown in a 7 cm cubic pot in our custom soil mix (30% peat moss, 10% perlite, 15% sand, 15% Sunshine potting soil, 20% vermiculite and 10% water) supplemented with Osmocote 14-14-14 (N-P-K) fertilizer at 3.3 g/L. The reactions of those lines to stripe rust were evaluated with the current predominant race PSTv-37, virulent to *Yr6*, *Yr7*, *Yr8*, *Yr9*, *Yr17*, *Yr27*, *Yr43*, *Yr44*, *YrTr1*, *YrExp2*, and is avirulent to *Yr1*, *Yr5*, *Yr10*, *Yr15*, *Yr24*, *Yr32*, *YrSP*, *Yr76.* The fresh urediniospores of PSTv-37 were mixed with Novec 7100 fluid (3M Inc., St. Paul, MN, USA) at a ratio of 10 mg/mL and evenly sprayed on wheat plants at 2-leaf stage with an atomizing sprayer. The plants were incubated at 10°C in a dark dew chamber for 24 hr, subsequently, the plants were moved to a controlled growth chamber with the temperature gradually increasing from 4°C at 4:00 am to 22°C at 4:00 pm then decreasing to 4°C with 16 hr light/8 hr dark for stripe rust development. The IT score of each EDR line was recorded 18 days after inoculation using a 0 to 9 IT scale (Line and Qayoum 1991) of which IT 0 to 3 was considered resistant, IT 4 to 6 was considered intermediate, and IT 7 to 9 was considered susceptible.

#### Leaf rust

##### *P. triticina* races MNPSD and MPPSD inoculation

Two seeds of each line were sown in a single cell of 72-celled trays filled with a commercial ‘Redi-Earth’ soil (Sun Gro, Bellevue, WA, USA) Plants were raised in a rust-free greenhouse at 20°C/18°C (day/night) and a 16 hr photoperiod. Jack’s classic® all-purpose (20-20-20) fertilizer was applied to the seedlings once a week. The EDR lines were planted in 4 replications. Leaf rust susceptible checks such as Thatcher, TAM110, Chisolm and Danne and a resistant check, Thatcher-*Lr19* were included in the test (Table S6). Seedlings were inoculated at the 2^nd^ leaf stage with a suspension of urediniospores comprised of a mixture of races collected from Oklahoma in 2021 and 2022 (primarily *P. triticina* races MNPSD and MPPSD) and light mineral oil (Soltrol 170; 1 mg of urediniospores per 1 mL of oil) using an inoculator pressurized by an air pump. The oil was allowed to evaporate from the leaves for at least 30 min before incubating in dark at 20°C for about 18 hr and relative humidity of 100%. Following incubation, plants were transferred to the greenhouse at 20°C/18°C (day/night) and a 16 hr photoperiod.

Leaf rust infection types (ITs) were assessed 12 days post inoculation on the first leaves of seedlings using a 0-to-4 scale(Stakman et al. 1962) where IT 0 = no visible symptom, ; = hypersensitive flecks, 1 = small uredinia with necrosis, 2 = small- to medium-size uredinia surrounded by chlorosis, 3 = medium-size uredinia with no chlorosis or necrosis, and 4 = large uredinia with no necrosis or chlorosis. [AM1] Larger and smaller uredinia than expected for each IT were designated with + and −, respectively. Lines showing ITs of 0 to 2+ were considered resistant, while 3 and 4 scores were considered susceptible(Long and Kolmer 1989; McIntosh et al. 1995). For data analysis, the 0-to-4 scale was converted to a linearized 0-to-9 scale as described by (Zhang et al. 2014a). Ratings of 0 to 6 were classified as resistant IT and 7 to 9 were considered as susceptible IT.

##### *P. triticina* race BBBQD inoculation

For the leaf rust screening, wheat seedlings at the two-leaf stage were evaluated for their reactions to BBBQD in the biosafety level 2 (BSL 2) facility at Dalrymple Research Greenhouse Complex, North Dakota Agricultural Experiment Station (AES), Fargo. Briefly, 3-4 seedlings per each EDR line along with susceptible checks were used for phenotypic screening for BBBQD with 3 technical replicates. Plants were grown in trays containing PRO-MIX LP-15 (Premier Tech Horticulture) sterilized soil mix and maintained in a rust-free greenhouse growth room set to 22°C/18°C (day/night) with 16 hr/8 hr light/dark photoperiod. At two-leaf stage, the seedlings were inoculated with fresh urediniospores suspended in SOLTROL-170 mineral oil (Phillips Petroleum) at a final concentration of 10^5^ spores per ml using an inoculator pressurized by an air pump. RL6089 and Little Club were used as the susceptible checks. The inoculated seedlings were placed in a dark dew chamber at 20°C overnight and then transferred back to the growth room.

The infection types (IT) were scored about 12 to 14 days after inoculation, using the 0-4 scale (Stakman et al. 1962). For each IT, ‘+’ or ‘-’ was used to represent variations from the predominant type. A ‘/’ was used for separating the heterogeneous IT scores between leaves with the most prevalent IT listed first. For plants with different ITs within leaves, a range of IT was recorded with the most predominant IT listed first. The IT scores were converted to a 0-9 linearized scale referred to as infection response (IR) (Zhang et al. 2014a). Plants with linearized IR scores of 0-4 were considered as highly resistant, 5-6 as moderately resistant, and 7-9 as susceptible.

#### Powdery mildew

Seeds of each line were sown in 8.25 in. × 1.5 in. diameter plastic cones containing a commercial ‘Ready-Earth’ soil (Sun Gro, Bellevue, WA, USA). The lines were planted in 4 replicates with two plants each. Plants were raised in the greenhouse at 20°C/18°C (day/night) and a 16 hr photoperiod. Seedlings were inoculated at the 2^nd^ leaf stage by gently shaking heavily infected plants of Jagger, infected with Oklahoma field isolates of *Blumeria graminis* f. sp. *tritici* over seedlings of the tested lines. Approximately 7-10 days after inoculation, infection types on the plants were rated on a 0-5 scale with 0 being the most resistant reaction and 5 being the most susceptible reaction. Plants having scores of 0-2 were considered resistant, plants having a score of 3 were considered intermediate, and plants having a score of 4 -5 were considered susceptible. Powdery mildew susceptible check variety Jagger was included as a control.

#### Tan spot

Seeds of each line were sown in 8.25 in. × 1.5 in. diameter plastic cones containing a commercial Redi-Earth’ soil (Sun Gro, Bellevue, WA, USA). The lines were planted in 8 reps with two plants in each rep. Plants were raised in a growth chamber at 20°C/18°C (day/night) with a 16 hr photoperiod. Tan spot susceptible checks ‘Billings’ and TAM 105, an intermediate check ‘Karl 92’, and a resistant check ‘Red Chief’ were included in the test. Wheat seedlings were inoculated at the third leaf stage (1^st^ and 2^nd^ leaves fully expanded) using an atomizer (DeVilbiss Co., Sommerset, PA, USA) with a conidial suspension (2,000 conidia per ml). An Oklahoma isolate of *P. tritici-repentis* Race1 was used for the inoculation. About one hour after inoculation, when conidia adhered to dried leaves, seedlings were placed in a mist chamber that provided near 100% relative humidity for 48 hr. Plants were then placed in a growth chamber at 20°C/18°C (day/night) with a 16 hr photoperiod. One week after inoculation, 1^st^ and 2^nd^ leaves were rated to determine percent leaf area infected by tan spot. For further analysis, we considered severity on the 2^nd^ leaves.

### Remapping EDR mutant exome capture data to new Kronos long-read assembly

The exome capture data for a non-mutagenized control (Kr0) and 32 EDR lines were downloaded from the National Center for Biotechnology Information (NCBI) and mapped to the Kronos reference genome (Seong et al. 2023 https://zenodo.org/records/10215402). Two EDR lines, Kr2027 and Kr2067, were excluded in this analysis as their data were unavailable. The paired-end datasets were first trimmed and filtered to remove sequencing adapters and low-quality reads with fastp v0.23.2 (Chen et al. 2018b). The processed reads were mapped to the Kronos genome with bwa aln v0.7.17-r1188 (Li 2013). The alignments were sorted with samtools v1.15.1 (Li et al. 2009), and duplicates were removed with picard v3.0.0 (https://github.com/broadinstitute/picard). These filtered alignments were processed with the MAPS pipeline (Henry et al. 2014). A minimum mapping quality of 20 was required for alignments to be processed. A minimum of 20 libraries was needed to identify valid mapping positions. HetMinPer was set to 15, and the following pairs of HomMC and HetMC were chosen: (2, 3), (3, 2), (3, 4), (3, 5), (4, 3), (5, 3) and (6, 4). Regions of residual genetic heterogeneity were removed. For each mutant individual library, the parameter producing a maximum number of mutations with a EMS-type mutation rate of 98% or higher was selected (Zhang et al. 2023).

### Mutation effect analysis

The mutation effect in each EDR line was examined with snpEff v5.2a (Cingolani et al. 2012). All analyses relied on a high-confidence gene set that contains 69,808 genes, unless specified. The effect was analyzed on a transcript per gene that encodes the longest protein sequences.

### Orthology inference

The putative orthology was inferred with OrthoFinder v2.5.5 (Emms and Kelly 2019). Proteomes from six species were analyzed, including *Arabidopsis thaliana* (Araport11) (Cheng et al. 2017), *Oryza sativa* (IRGSP-1.0) (Kawahara et al. 2013), *Solanum lycopersicum* (ITAG 4.0) (Hosmani et al. 2019), *Triticum aestivum* (IWGSC) (International Wheat Genome Sequencing Consortium (IWGSC) 2018), *Zea mays* (NAM 5.0) (Woodhouse et al. 2021) and Kronos. Both high and low-confidence gene models were included for Kronos. Only the longest protein products per gene were used. OrthoFinder relied on Diamond v2.1.7 (Buchfink et al. 2015) for homology inference, MAFFT v7.525 (Katoh et al. 2002) for sequence alignments and FastTree v2.1.11 (Price et al. 2010) for constructing phylogenetic trees. Orthogroups that included known LMMs, suppressors of LMMs and Yr gene products were identified. Orthology was categorized as one-to-many, many-to-one or many-to-many, based on the number of proteins from a reference species and Kronos. Typically, only a limited number of members were identified in most analyzed orthogroups for Kronos and common wheat. In more complex cases involving gene duplications and deletions, reciprocal best blast searches were employed to match putative orthologs between two species (Camacho et al. 2009). For Yr gene products, phylogenetic trees were also utilized to infer orthology.

### Statistical analysis

Figures and graphs were generated using GraphPad Prism and at BioRender.com. Means were compared using ordinary one-way analysis of variance and Fisher’s LSD test or unpaired t-tests, for the quantitative RT-PCR comparisons.

## Discussions

In this work, we described 34 independent EMS-mutagenized Kronos wheat lines with increased adult plant resistance to stripe rust. After initial identification, we tested the consistency of the resistance by planting the candidate lines over several growing seasons. During our observations in a fungicide-treated field, we noticed that several mutant lines had additional ‘flecking necrosis’ or lesion mimic phenotypes even without the pathogen, indicating autoimmunity. In growth chamber experiments, we found this phenotype to be enhanced in cooler temperatures. We also found a differential response to other fungal pathogens at both the seedling and adult plant stages. To test for defense-growth trade-offs in our stripe resistant lines, we measured kernel weights under pathogen infection and found most EDR lines to be the same or better than Kronos. The EDR lines described herein are likely to contain valuable EMS-generated mutations to uncover both mechanisms of plant-pathogen interaction and for applied breeding programs.

Several categories of stripe rust resistance have been identified and these differ in the stage of the plant in which they are active, their modes of inheritance and durability. Combining genes that confer resistance at different stages and by different mechanisms is desirable to combat diverse pathogen populations (Wang and Chen 2017). The EDR collection described here should complement previously identified enhanced resistance durum wheat germplasm as it has diverse resistance phenotypes. Genes conferring resistance to microbial pests that also cause lesion mimic phenotypes are known across many plant systems (Bruggeman et al. 2015). Many mutants with lesion mimic phenotypes also confer resistance to pathogens and several of these have been cloned in crops, including *Mlo* from barley, (Büschges et al. 1997), *Spl26* (Ting et al. 2019) and *Spl7* (Yamanouchi et al. 2002) from rice, and *Rp1* (Hu et al. 1996; Hulbert 1997) from maize. Interestingly, some maize *Rp1* alleles are autoimmune, but other alleles require pathogen stimulus to cause cell death (Hu et al. 1996). The *Rp1-D21* allele is autoimmune with a differential response to temperature; lesions are suppressed at high temperatures and enhanced at lower temperatures (Negeri et al. 2013). Lesion mimic mutations that also provide disease resistance may be useful targets for gene editing. Recently, by combining forward lesion mimic phenotype screens of a mutagenized rice population with gene-editing in rice, a broad-spectrum loss of function allele of a susceptibility gene, *RESISTANCE TO BLAST1 (RBL1)*, encoding a phospholipid biosynthesis gene, was created giving resistance to both *Magnaporthe oryzae*, the causal fungal pathogen of rice blast and *Xanthomonas oryzae* pv. *oryzae* which causes rice bacterial blight (Sha et al. 2023). Gene-editing will aid in fine-tuning and engineering pathogen resistance and forward screens are an efficient method to identify target sequences for this editing.

We sought to further characterize the EDR lines in reaction to diverse pathogens, in search of lines that may have resistance to multiple diseases caused by pathogens with different lifestyles. The stripe rust pathogen, *P. striiformis*, is an obligate biotroph (Ralf T. Voegele et al. 2009) requiring living tissue to sustain its growth. In contrast, *Z. tritici* and *P. tritici-repentis* gain entry to host tissues by causing cell death and colonizing necrotic tissues after a variable latent period which may or may not be biotrophic (Steinberg 2015). Given this difference in lifestyle, we hypothesized that EDR lines, which utilize cell death constitutively due to autoimmune triggers or early during the hypersensitive response (HR) to restrict biotrophic pathogens, are at an increased risk of colonization by necrotrophic pathogens (Kliebenstein and Rowe 2008).

Nearly a third of the persistently stripe rust resistant lines we identified were autoimmune, suggesting that these genes could provide broad spectrum, durable resistance. However, the autoimmune lines tended to fare slightly worse than the other EDR lines under challenge by the latently necrotrophic pathogen, *Z. tritici*, and the necrotrophic pathogen, *P. tritici-repentis*, suggesting a biotrophic-necrotrophic trade-off, as has been previously reported. In oat, most notably, HR induced by the *Puccinia coronata* resistance gene *Pc2* is purported to be allelic to the *Vb* gene that conditions susceptibility to Victoria blight, caused by the necrotroph, *Cochliobolus victoriae* (Lorang et al. 2007, 2010, 2012). In dicots, salicylic acid (SA) and jasmonic acid (JA) signaling are antagonistic; SA is induced in response to biotrophs and JA is induced in response to necrotrophs and herbivores (Stout et al. 2006; Kliebenstein and Rowe 2008). However, in rice, which has endogenously high levels of SA, both hormone pathways converge to confer immunity through a common pathway that can be activated by either hormone and can confer resistance to pathogens irregardless of lifestyle. This was uncovered by microarray studies that showed many genes were upregulated by both JA and an SA analog (Tamaoki et al. 2013). Broadly, perturbation of many pathways can cause spontaneous lesions: immunity mediated by NLR resistance genes, metabolism of fatty acids, hormones, photosynthesis, reactive oxygen species production and programmed cell death (Zhang et al. 2022a) a process that is only partially understood (Maekawa et al. 2023). The availability of these newly described durum lesion mimic lines provides a resource for further study of mechanisms of plant cell death in wheat. Further experiments could characterize these mutants by the morphological aspects of their cell death phenotypes (van Doorn et al. 2011). We found that Pr1, Pr3 and ZLP are induced in 2 of the 3 autoactive EDR lines we tested and seemed correlated with the severity of the lesion phenotype. This suggests at least in a subset of EDR lines the LMM phenotype is consistent with ectopic activation of known plant immune pathways. Further characterization of these mutants by transcriptomic and metabolomic analyses as well as identification of causative alleles would uncover lesion mimic pathways shared across plants.

The EDR lines that had autoimmune lesions did perform more poorly in tests with necrotrophic pathogens but they did not have a reduction in TKW. Resistance versus yield tradeoffs have been previously documented (Draz et al. 2015; Olukolu et al. 2016) and a more comprehensive yield analysis should be done after the EDR lines have been incorporated into breeding programs. Susceptibility of EDR lesion mimics to necrotrophs was milder at the seedling stages, as expected, as visible, external lesions only appear after the juvenile phase. Additionally, since the EDR lines were phenotypically selected for high-temperature adult-plant resistance, it is possible that some lines carry all-stage resistance and that many solely carry adult resistance. Although the FHB pathogen, *F. graminearum*, is also a mostly necrotrophic pathogen (Kheiri et al. 2019), the autoimmune EDR lines performed similarly to Kronos in our trials. The EDR lines did show a differential response to FHB as was seen in an EMS population of hexaploid wheat, ‘Jagger’ (Chhabra et al. 2021a). We surmise that the lack of correlation between stripe rust resistance and FHB resistance may be because FHB is a disease of the inflorescence rather than a foliar disease. Our phenotypic selection for the foliar stripe rust disease may have favored leaf-specific resistance mechanisms.

In the stripe rust field trials, we did not see a penalty as measured by TKW, so phenotypic selection for resistance at the adult stage is an effective approach to find useful resistance alleles with favorable agronomic traits. However, TKW is only one yield component and a comprehensive evaluation of yield performance of these lines in fields with low and high pathogen pressure, once they have been combined with additional improved germplasm, should be explored. The allele combinations in the EDR lines provide a valuable resource for researchers exploring multiple aspects of plant immunity. Almost a third of the mutants are lesion mimics, a class of mutants which has provided extensive information about programmed cell death. Overall, we show that our identified EMS lines in an elite durum wheat have enhanced resistance to stripe rust and other fungal pathogens. The mutations in these lines have already been sequenced and incorporated into genome browsers and include candidate mutations in previously known EDR genes. Our study includes remapping of these mutations to a recent high-quality Kronos genome, and we expect this resource will greatly aid in the identification of causal variants, and their deployment into both basic and applied research programs.

## Supporting information

Supplemental figures all

Supplemental tables all

## Data availability

The mutant mapping data produced in this study are available in Github: https://github.com/krasileva-group/Kronos_EDR. See Table S9a for NCBI accession numbers of exome data. The new Kronos assembly and annotation v1.0 is available at https://zenodo.org/records/11106422. The data generated in the re-analysis of exome capture data can be accessed through https://zenodo.org/records/11099763.

## Acknowledgements

We would like to thank Drs. Jorge Dubcovsky and Joshua Hegarty for their help in the forward screen and for sharing field space and resources. Francine Paraiso was also instrumental in the technical execution of the forward screen. Thank you to Drs. Robert Van Steenwyk of UC Berkeley and Derrick Hammons from Syngenta who advised us on the use of and procured Cruiser Maxx Vibrance seed treatment for the STB trials. Thank you to Duncan Ball for providing temperatures in Norwich, UK © Crown Copyright [2015]. Information provided by the National Meteorological Library and Archive – Met Office, UK. We also are grateful for the greenhouse and growth chamber support provided by Christina Wistrom and the staff at the Oxford Tract Greenhouse Complex at UC Berkeley. Thank you also to members of the Krasileva lab especially Dr. Wei Wei and Chandler Sutherland for helpful comments on the manuscript.

## Statements & Declarations

### Author Contributions

Material preparation, data collection and analysis were performed by China Lunde, Kyungyong Seong, Rakesh Kumar, Andrew Deatker, Bhavit Chhabra, Meinan Wang, Shivreet Kaur, Sarah Song, Ann Palayur, Cole Davies, William Cumberlich, Upinder Gill, Nidhi Rawat, Xianming Chen, Meriem Aoun, Christopher Mundt, and Ksenia V Krasileva. The first draft of the manuscript was written by China Lunde and Kyungyong Seong and all authors commented on previous versions of the manuscript. All authors read and approved the final manuscript.

