## Supplemental figures all for "Wheat Enhanced Disease Resistance EMS-Mutants Include Lesion-mimics With Adult Plant Resistance to Stripe Rust"

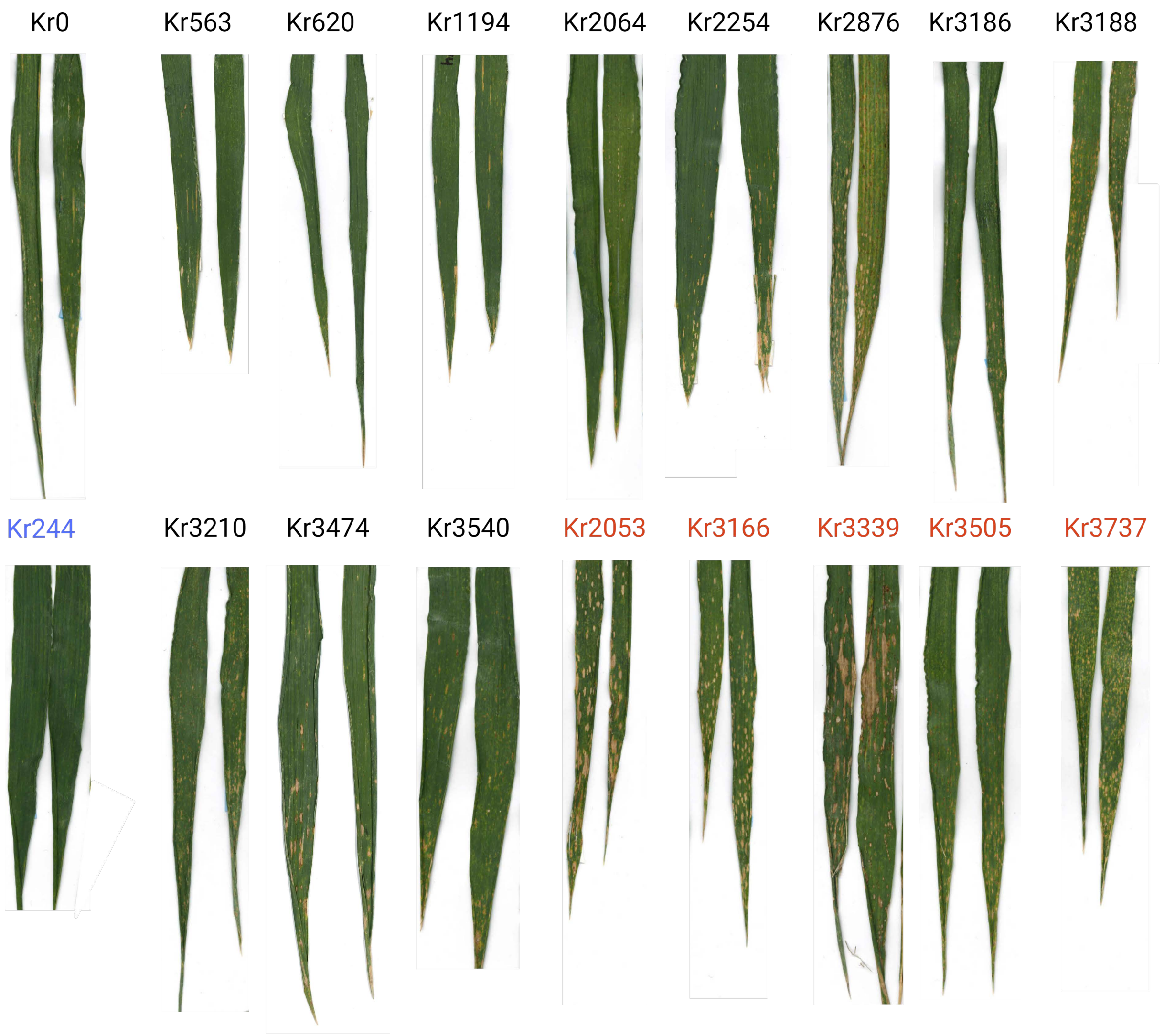


**Figure S1a.** Leaf scans of flag leaves of EDR lines with persistent stripe rust resistance over four years collected from plants grown in an inoculated field in Davis, CA using spores collected the previous season. These collected in April, 2020. EMS progenitor line Kronos (top row left) and EDR line Kr244 (bottom row, left, blue) with no visible cell death are included for comparison with the autoactive lesioned lines (red).

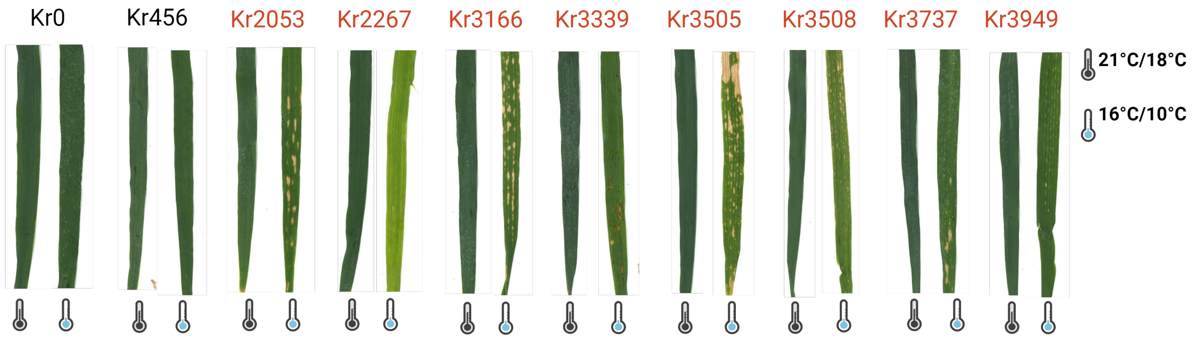


**Figure S1b.** Side by side leaf scans of flag leaves of putative autoimmune lesion mutants Eight of the nine identified lines had lesions under these conditions, but none had lesions under standard greenhouse conditions (21°C/18°C day/night (+/- 2°C) 16/8 hr). Kr456 did not form lesions under these conditions.

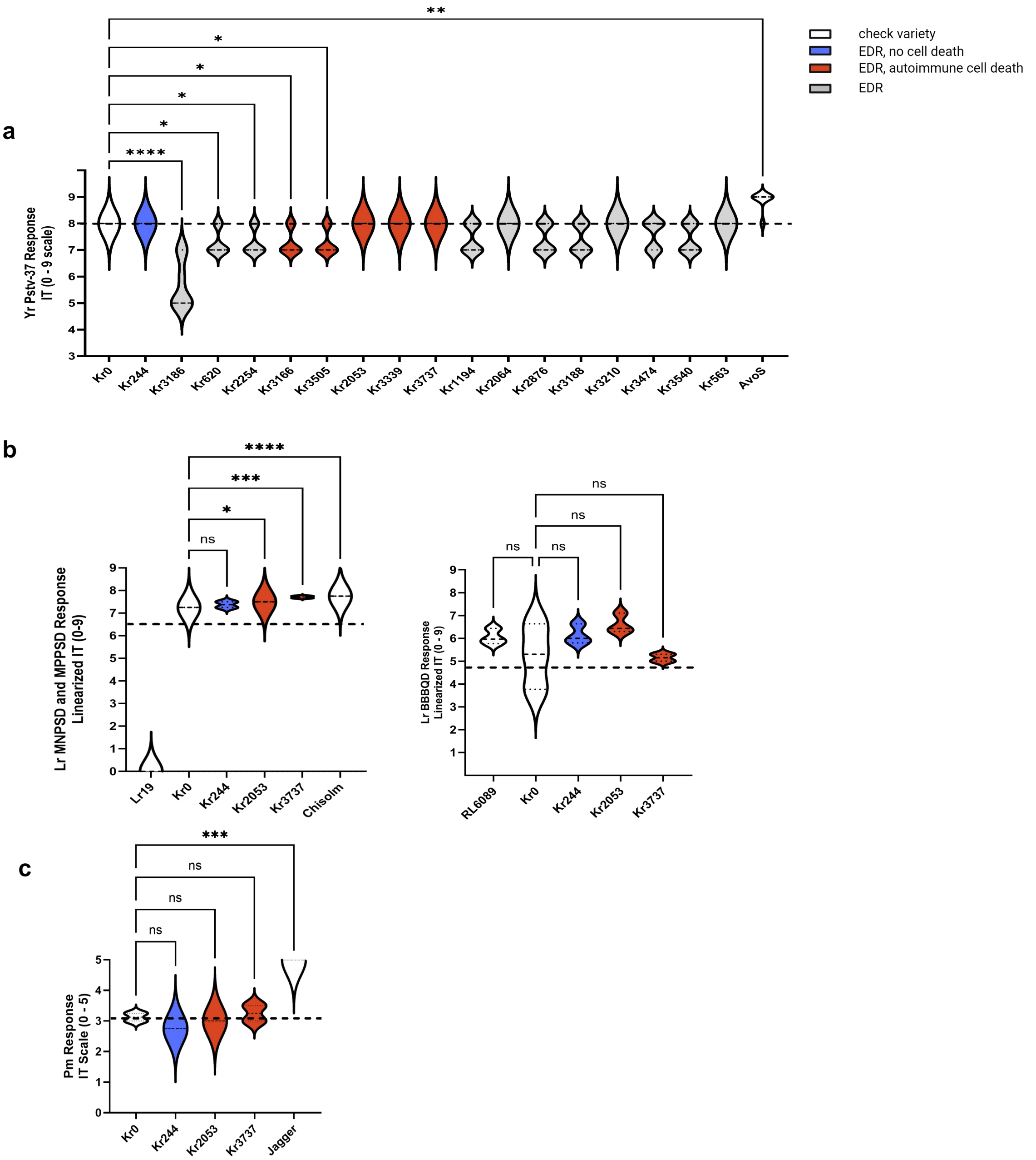


**Figure S2.** EDR seedling response to biotrophic pathogens: stripe rust, leaf rust and powdery mildew under controlled conditions A. Response of the 16 persistently resistant EDR lines to stripe rust Pstv-37. Lines indicated by asterisks have significantly lower ITs than Kronos. AvoS susceptible check has a significantly higher IT. B. Response of select lines to two Oklahoma leaf rust isolates (left, including resistant check Lr19 and susceptible check Chisolm) and LrBBBQD (right, including resistant check RL6089). C. Response of select EDR lines and susceptible check Jagger to powdery mildew. Control lines Kronos and other checks (white) plus an EDR line with no macroscopic cell death under pathogen challenge (blue) are included for comparison with the autoactive lesioned lines (red). Means were compared using one-way ANOVA, *, significant at $p=0.05$; **, significant at $p=0.01$; *** significant at $p=0.001$; ****, significant at $p<0.0001$; ns, non-significant.

**a**

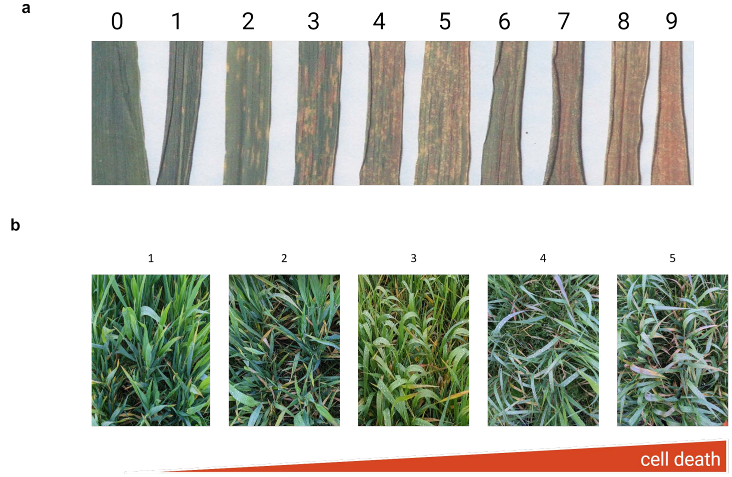


**Figure S3.** Stripe rust and STB field scoring rubrics a) Example flag leaves of each McNeal (1971) from 0 (no infection) to 9 (leaves completely covered in spores) collected in Davis, CA, USA in April, 2023. b) Plots showing representative STB scores for subjective observations. Scale ranges from 1 (little or no infection) to 5 (all plants showing chlorosis, especially of the lower canopy).

**Figure S4.** Response of EDR lines to STB
Five pages (to follow) of scanned leaves collected from STB field in April of 2022, in Corvallis, OR, USA

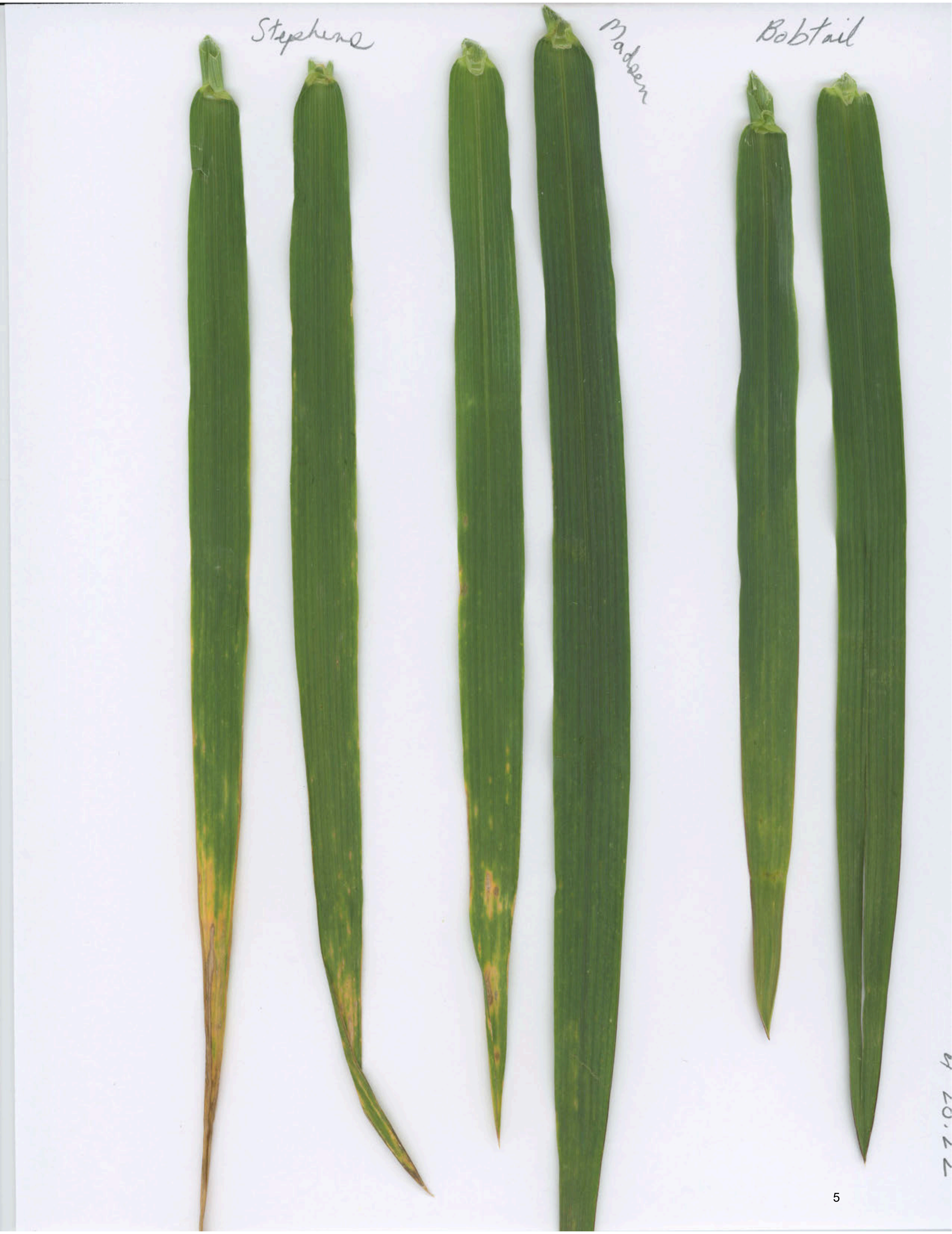

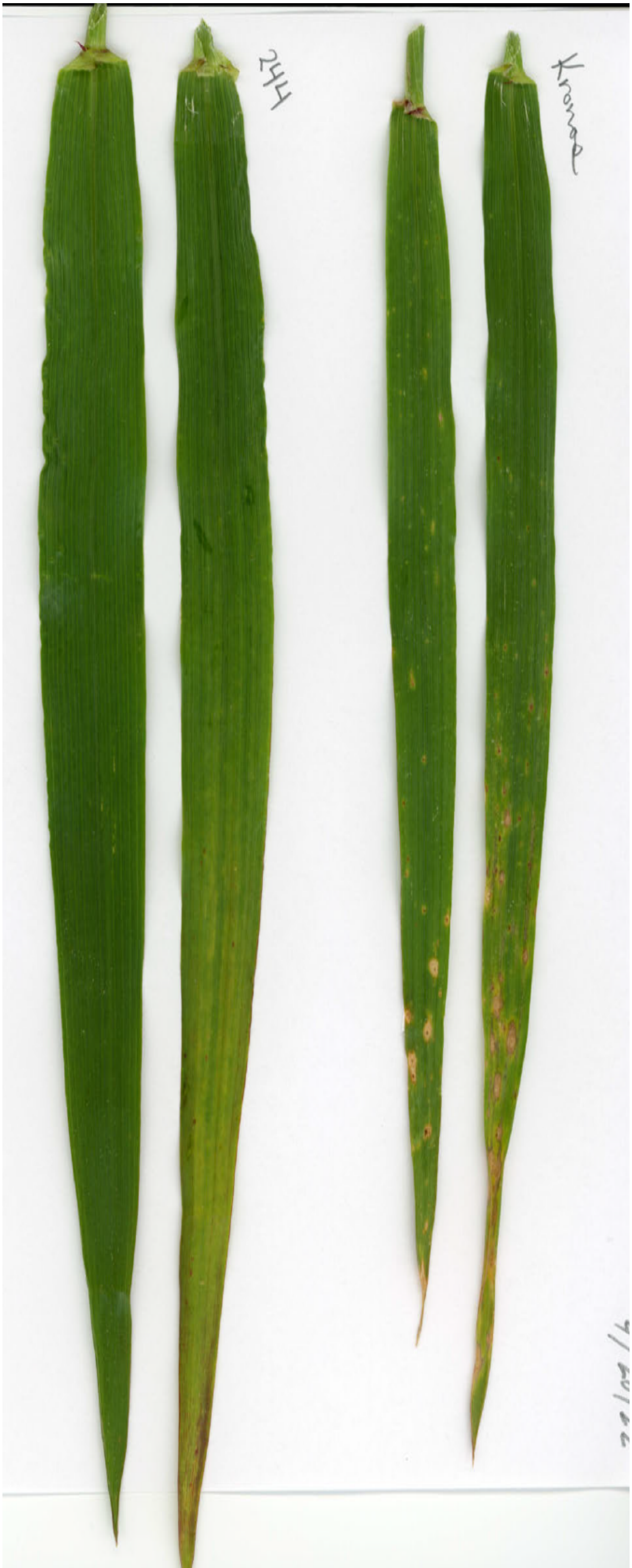

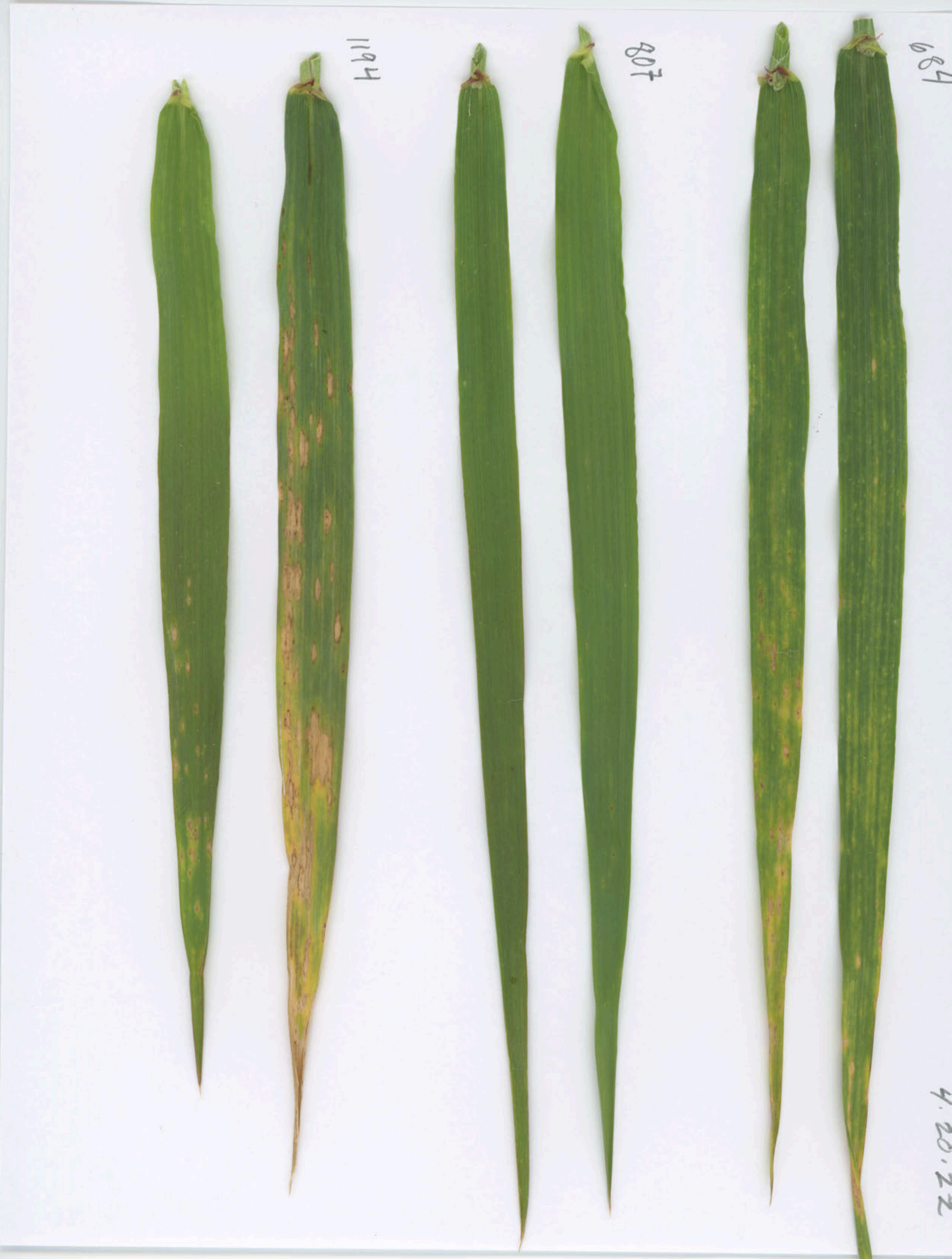
1194
807
4.20.22

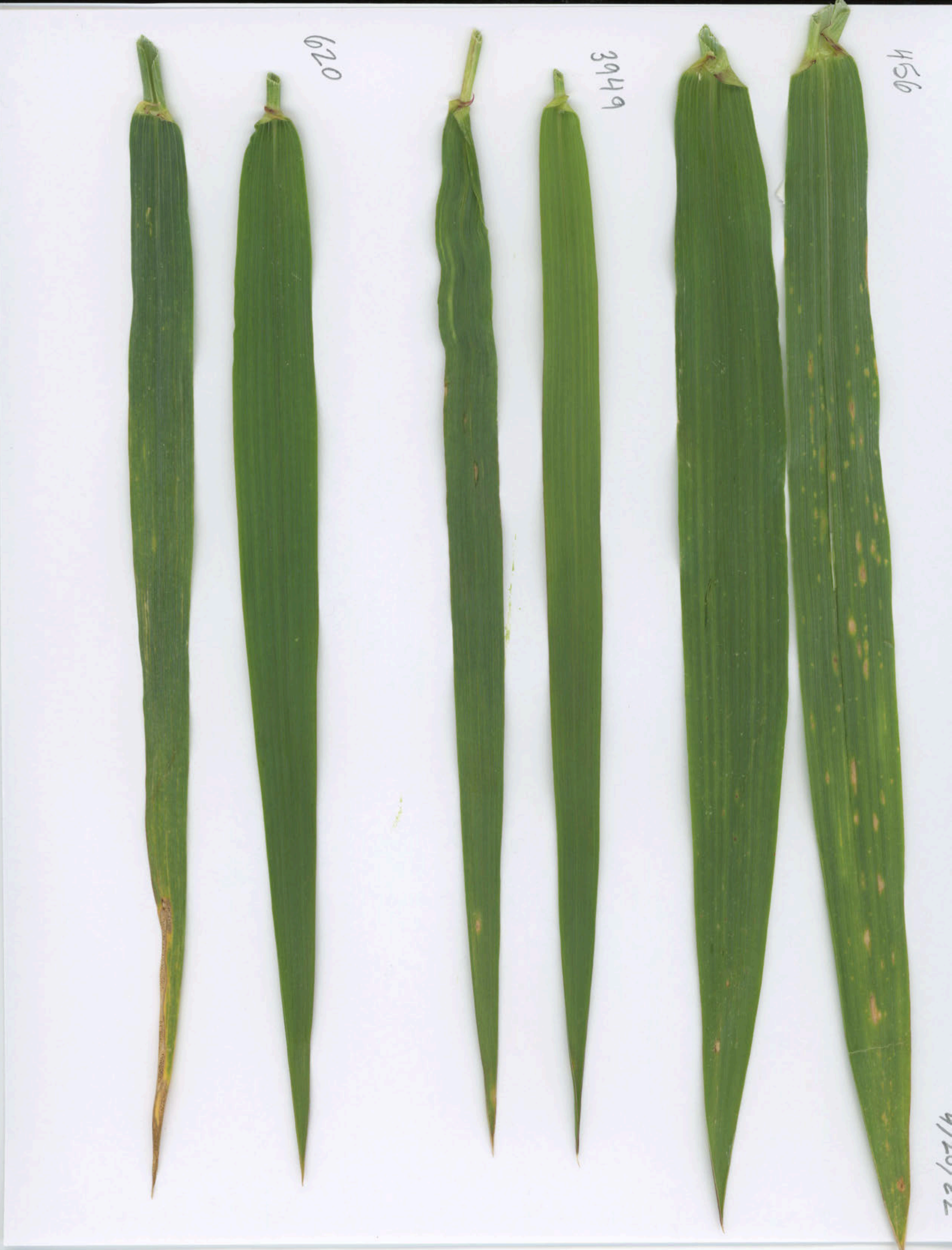
620
3949
456

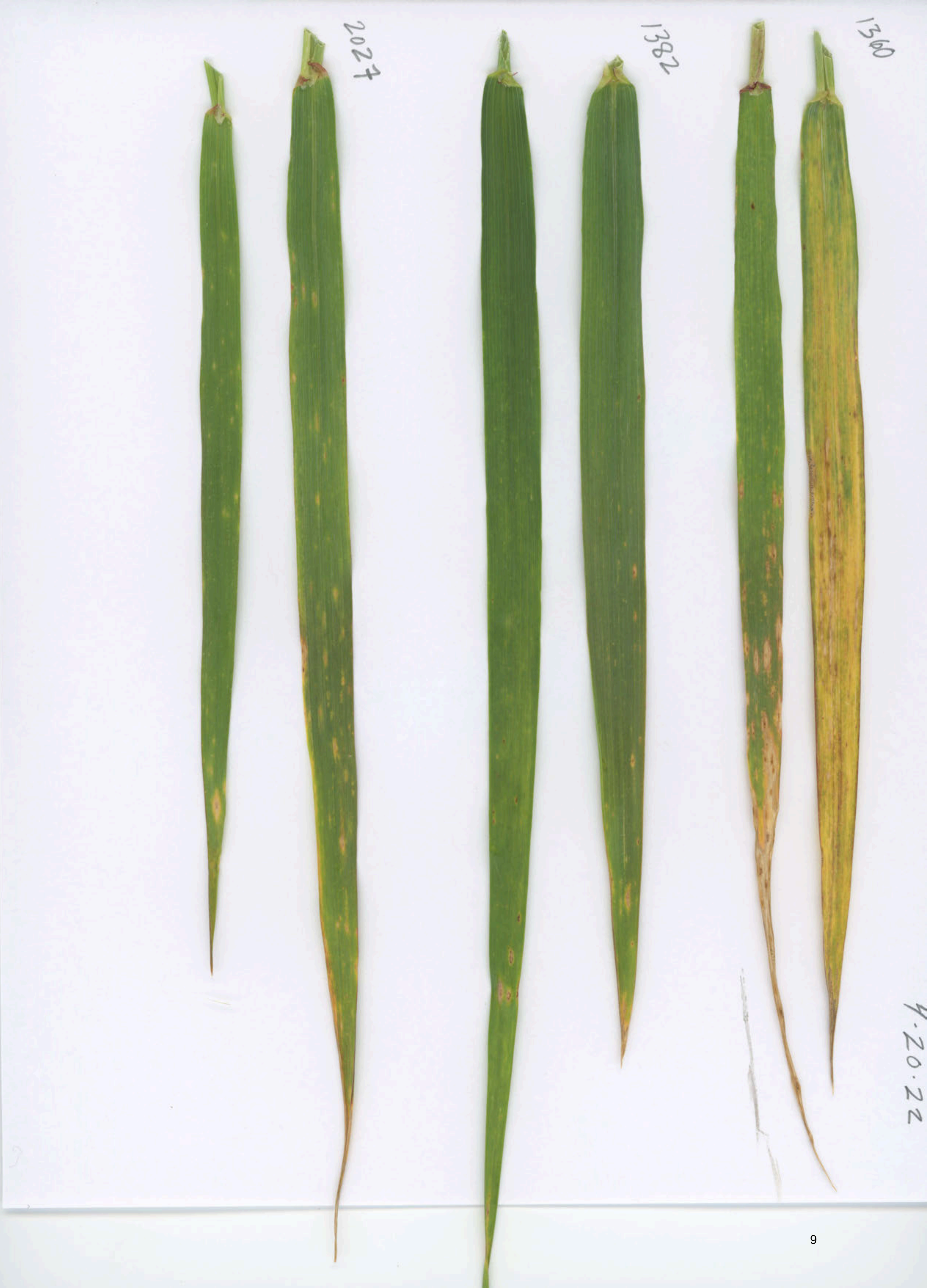
2027
1382
1360
4.20.22

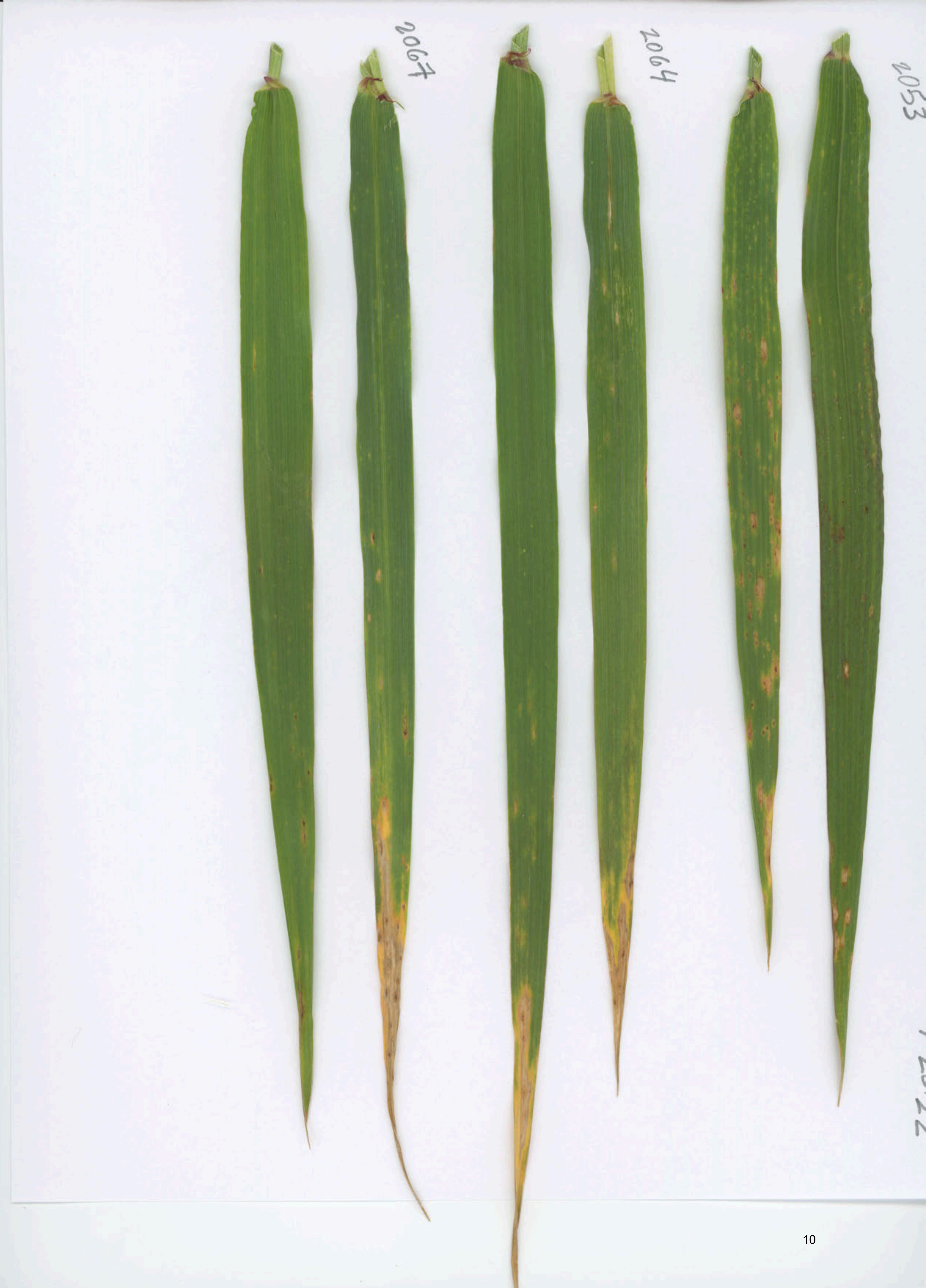
2067
2064
2053

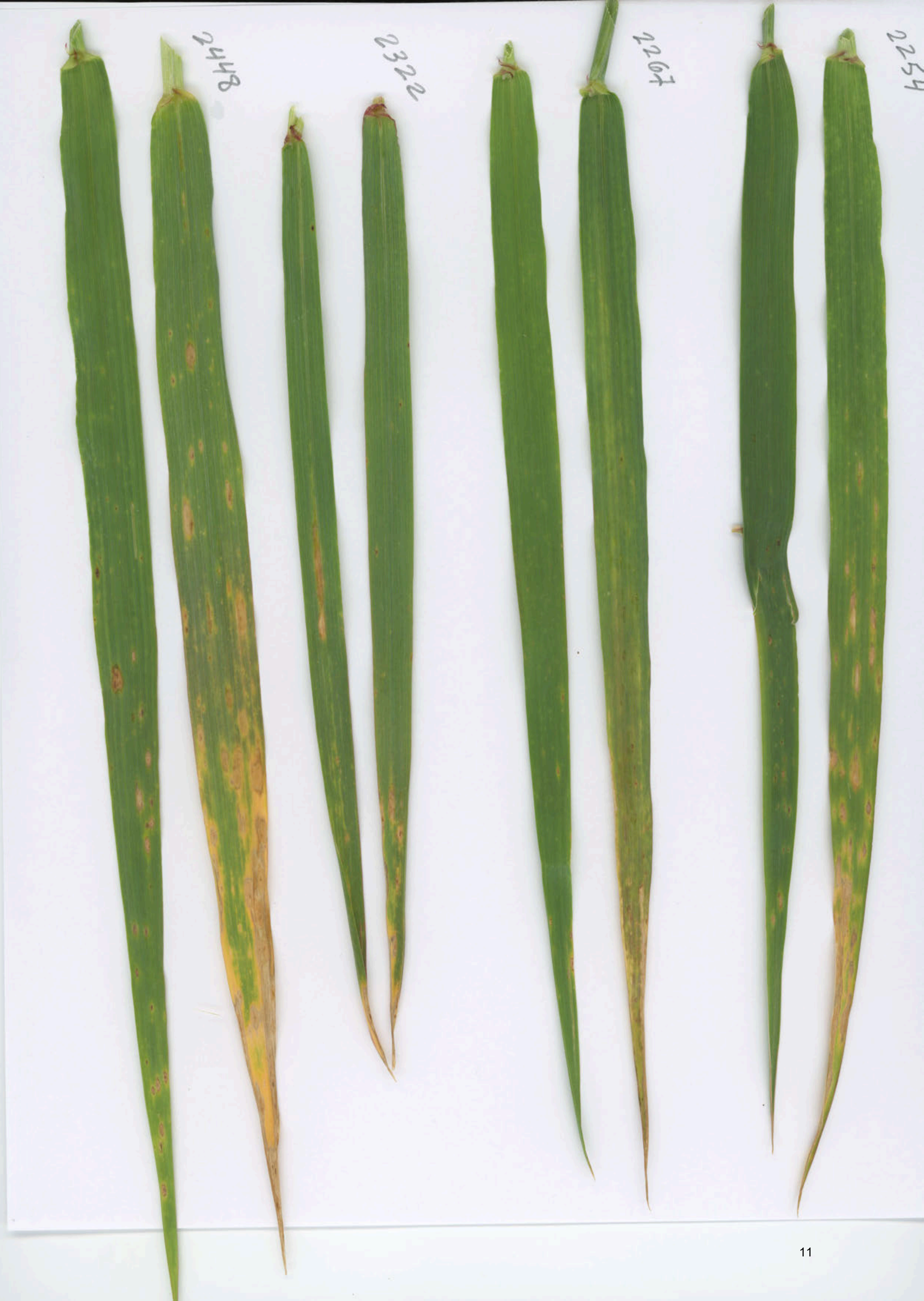
2448
2322
2267
2254

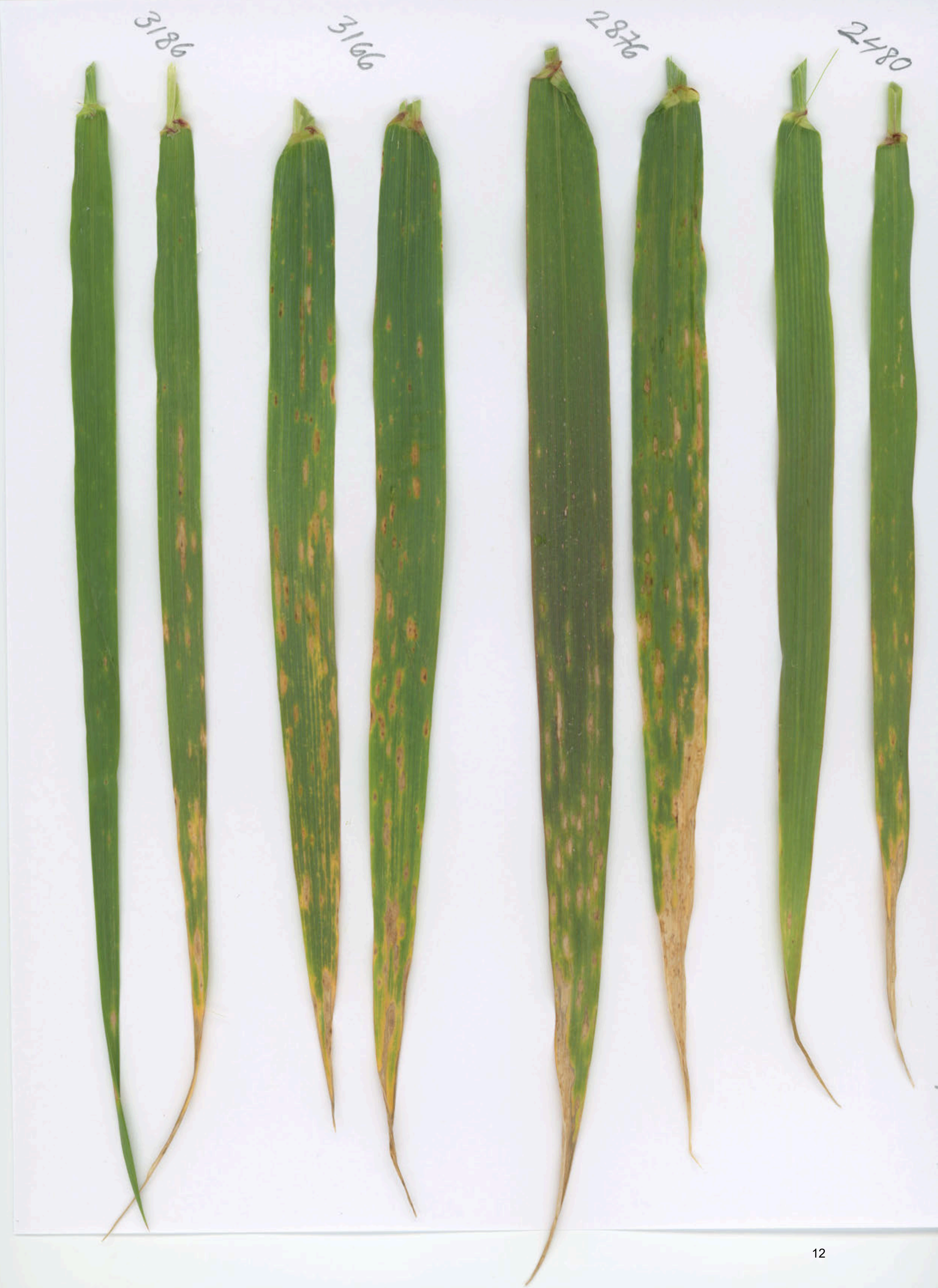
3186
3166
2876
2480

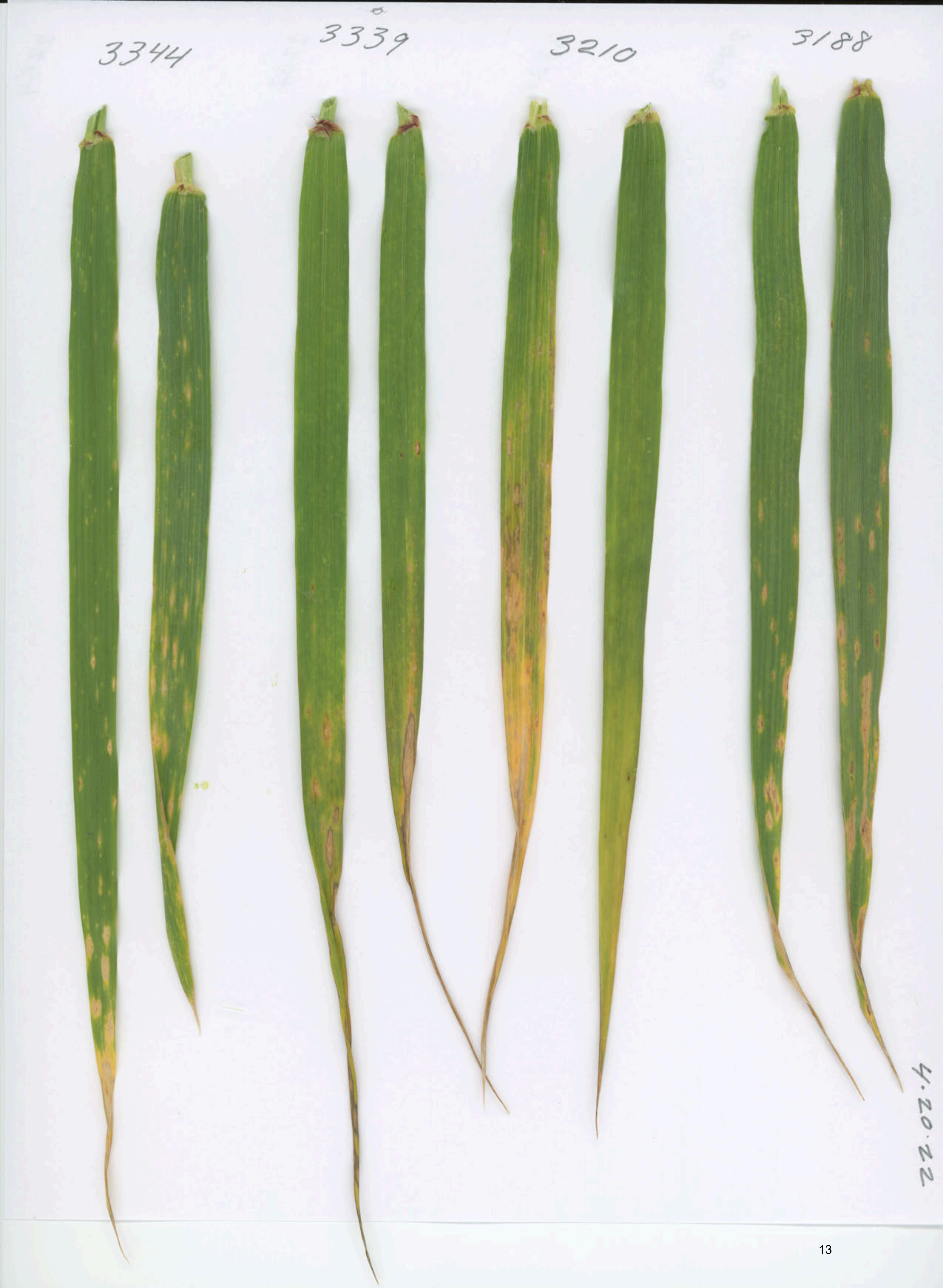
3344
3339
3210
3188
4.20.22

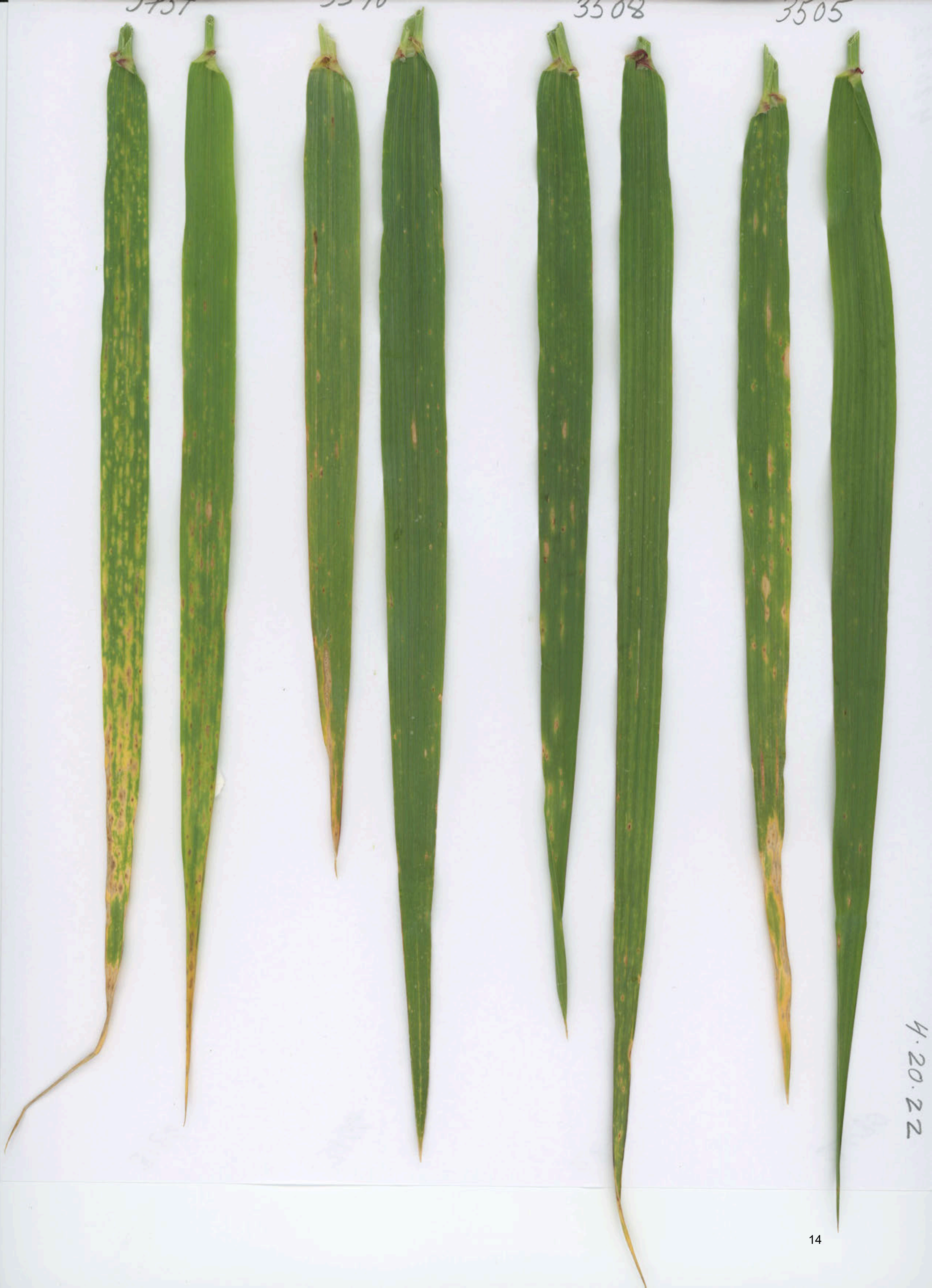
3508
3505
4.20.22

3474

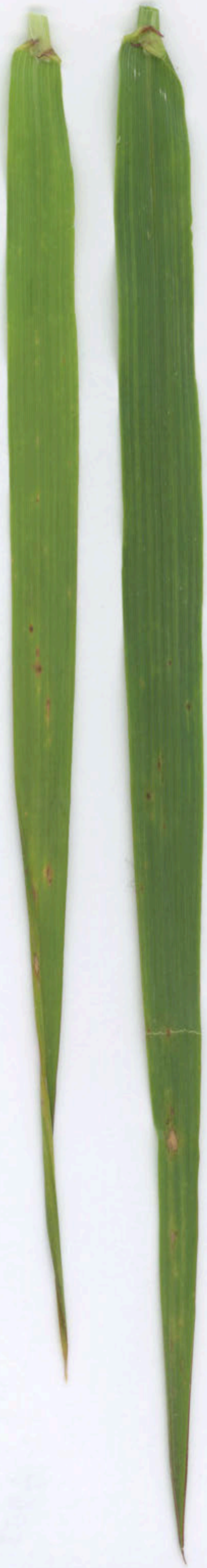

4.02.22

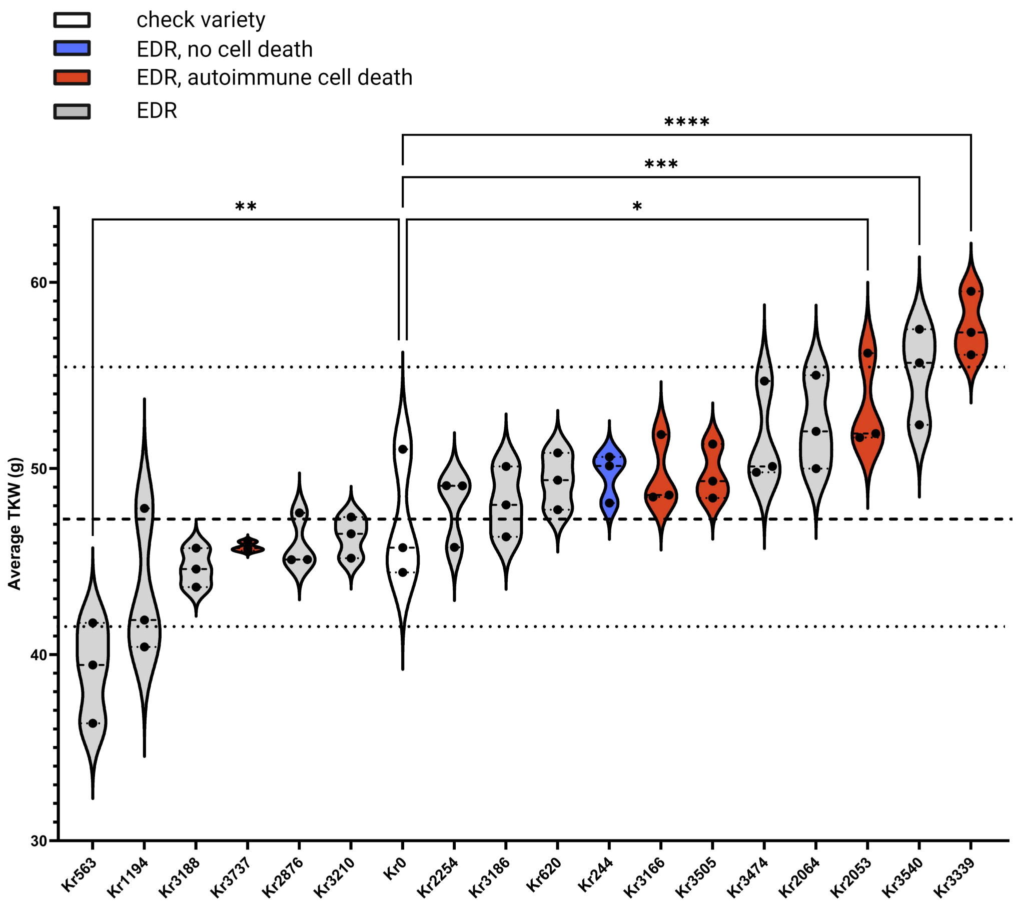


**Figure S5.** EDR lines do not show a TKW penalty when grown in stripe rust fields
Ranked mean TKW of EDR lines harvested in 2020 to 2022 in Davis, CA. Each point represents a year of data. Control line Kronos (white) plus an EDR line with no macroscopic cell death under pathogen challenge (blue) are included for comparison with the autoactive lesioned lines (red). Dotted line indicates Kronos median value. Means were compared using one-way ANOVA, *, significant at p=0.05; **, significant at p=0.01; *** significant at p=0.001; ****, significant at p<0.0001; ns, non-significant.

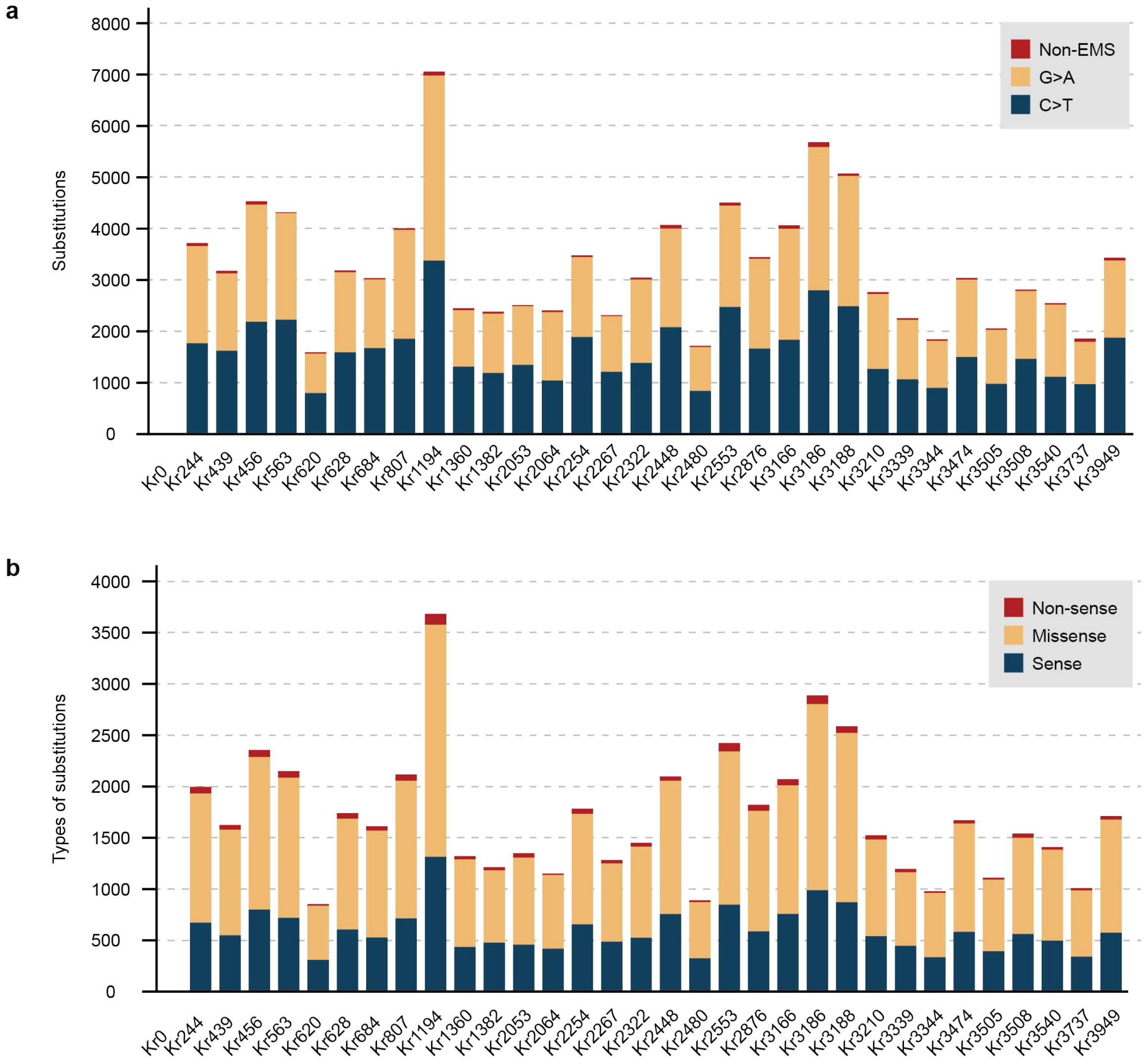


**Figure S6.** Identified substitutions in EDR lines and their impacts on proteins a) The number of substitutions identified with the MAPS pipeline from EDR lines and non-mutagenized control line Kr0. The mutations are colored to indicate non-EMS type mutations, and two EMS-type mutations (G to A and C to T). b) The mutation effects of identified substitutions in coding sequences.

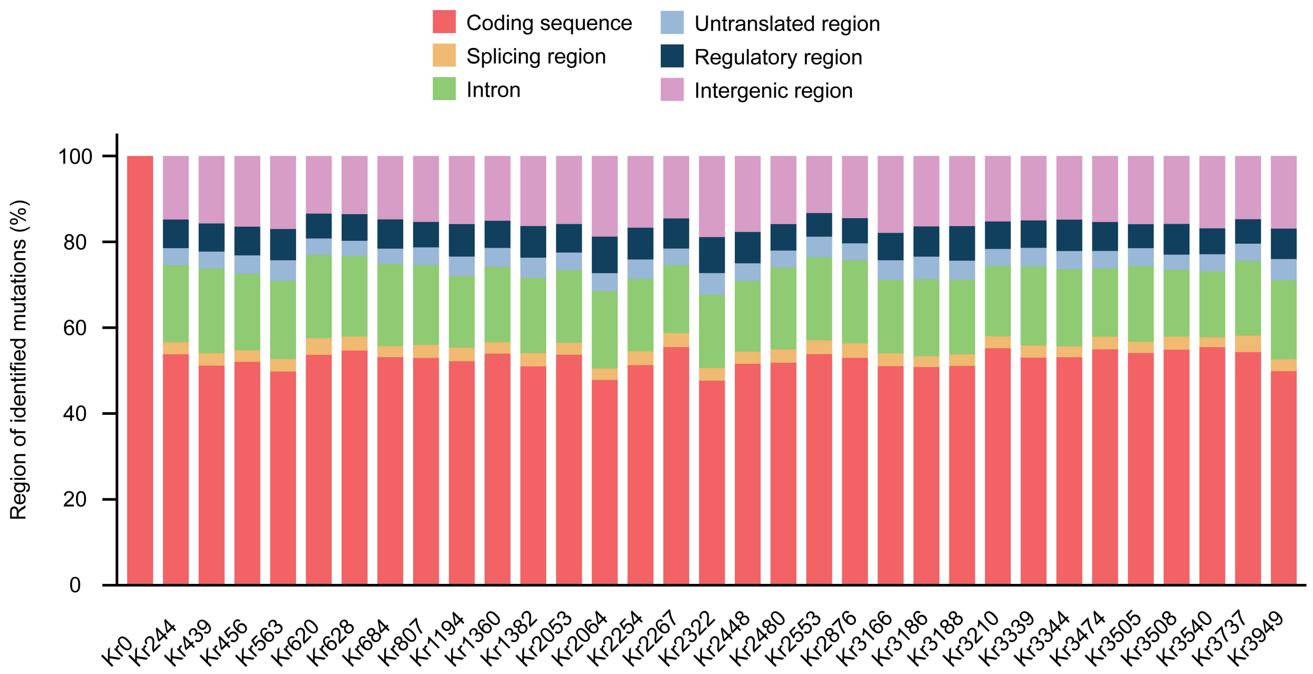


**Figure S7.** The classification of identified substitutions. Mutations identified in the EDR line were classified based on their location into the following categories: coding sequences, splicing regions, introns, untranslated regions, regulatory regions, and intergenic regions. Each mutation was counted only once, following the sequence of these classes.
